# Towards Identifying a Molecular Activator of Spreading Depolarization Generated by the Ischemic Brain

**DOI:** 10.64898/2026.04.27.721086

**Authors:** Chloe A. Lowry, Julia A. Hellas, Nikita Ollen-Bittle, Peter J. Gagolewicz, Brian M. Bennett, R. David Andrew

## Abstract

Spreading depolarizations (SDs) are waves of mass depolarization that propagate through gray matter following Na^+^/K^+^-ATPase (NKA) failure because of stroke, traumatic brain injury or sudden cardiac arrest. SDs expand the initial site of neuronal injury and worsen clinical outcomes. The molecular events underlying SD initiation and propagation are not well understood. In this rodent study, we hypothesized that gray matter stressed by oxygen/glucose deprivation (OGD) releases a compound(s) that promotes SD, which we term a spreading depolarization activator (SD*a*). We used rat brain slices incubated in artificial cerebrospinal fluid (aCSF) and subjected to OGD to release a putative SDa. The aCSF was collected either prior to (“Pre-SD aCSF”) or 10 min after initiation of OGD conditions (“Post-SD_OGD_ aCSF”). These solutions were then separately superfused over a healthy, naïve (non-stressed) brain slice. Post-SD_OGD_ aCSF (with re-normalized O_2_ and glucose) evoked SD in 82.35% of the naïve brain slices (*n* = 17) whereas Pre-SD aCSF evoked no SD in 10 naïve slices. Then to investigate the NKA as a potential target of the SD*a*, we used a hemolysis assay, comparing the effects of Pre- or Post-SD_OGD_ aCSF on red blood cell (RBC) lysis and compared it to the known hemolytic effect of the NKA-specific inhibitor, palytoxin.

Post-SD_OGD_ aCSF evoked neither swelling nor lysis of RBCs on its own. However, when a sub-threshold concentration (0.01–0.02 nM) of the specific NKA inhibitor palytoxin (PLTX) was added, a striking “priming” effect was observed, whereby Post-SD_OGD_ aCSF evoked a highly significant increase in both RBC swelling and then hemolysis, compared to Pre-SD aCSF. High pressure liquid chromatography (HPLC) experiments show a several-fold increase in released molecules post-SD vs pre-SD. Overall, this study provides support for SD*a* release capable of inducing SD-associated swelling in brain slices and, when combined with a trace amount of PLTX, swelling/hemolysis of RBCs caused by NKA inhibition. A greater understanding of the molecular events underlying SD should identify novel targets to reduce recurrent SD-evoked neuronal injury under ischemic conditions.

## Introduction

Within minutes of severe brain ischemia, depletion of available adenosine triphosphate (ATP) leads to slowing and eventual failure of the Na^+^/K^+^-ATPase (NKA) and subsequent collapse of transmembrane ion gradients necessary for proper neuronal function (Dreier, 2011; Hellas & Andrew, 2021). Spreading depolarizations (SDs) are waves of mass cellular depolarization that occur after NKA failure and that propagate through higher gray matter at a rate of 1–9 mm/min (Andrew, Hartings et al., 2022). SD arising from severe pathologies such as stroke, sudden cardiac arrest, or traumatic brain injury add to the energy demand within gray matter (Dreier & Reiffurth, 2015; Hartings et al., 2017; von Bornstädt et al., 2016). SD imposes a further bioenergetic burden upon ischemic tissue through a sudden loss of ionic gradients and resting membrane potential, which must be restored for neuronal survival and functional recovery (Andrew, Farkas et al., 2022; Andrew, Hartings et al., 2022; Pietrobon & Moskowitz, 2014). Furthermore, SD can elicit recurring vasoconstriction in the hours and days following brain injury, expanding the initial site of injury and worsening clinical outcome (Chung et al., 2016; Dohmen et al., 2008; Dreier et al., 2006; Strong et al., 2002). Importantly, the recurrent nature of SD provides a therapeutic window lasting many hours whereby SD events could be inhibited to mitigate the impending brain damage and improve functional outcomes (Andrew, Hartings et al., 2022; Hartings et al., 2003).

Although the NKA is a major player in ischemia-evoked SD, the molecular mechanism connecting pump failure to SD initiation is unclear (Andrew & Hartings et al., 2022; Hartings et al., 2003; Ricks et al., 2025; Withers et al., 2025). In addition, the channel(s) underlying the inward cationic macro-current that drives SD has yet to be identified, despite its predicted existence based on computer modelling (Makarova et al., 2007) and evidence that it is a non-specific Na^+^ /K^+^ conductance (Czéh et al., 1993; Czeh et al., 1992; Kim et al., 2025; Wang et al., 2024).

Further evidence of a novel channel driving SD is the difficulty in suppressing SD either *in vivo* or in live brain slices by standard antagonists of voltage-gated or ligand-gated ion channels (Aitken et al., 1991; Jarvis et al., 2001; Jing et al., 1993; Müller, 2000; Obeidat et al., 2000; Andrew, Farkaset al., 2022), including *N*-methyl-D-aspartate (NMDA) and α-amino-3-hydroxy-5-methyl-4-isoxazolepropionic acid (AMPA) receptors. Rather, SD could be driven by an unidentified channel that is normally silent in healthy brain tissue. For example, the conversion of an existing membrane transporter into an open channel during periods of metabolic stress (Kim et al., 2025; Wang et al., 2024; Andrew & Hartings et al., 2022). This concept has precedence from an evolutionary perspective whereby transporter and channel mutations can convert one into the other (Ashcroft et al., 2009; DeFelice & Goswami, 2007; Durell et al., 1999; Higgins, 1992; Miller, 2006). Some biological toxins exploit such underlying vulnerabilities. For example, the marine poison palytoxin (PLTX) converts the NKA into an open Na^+^/K^+^ channel in both excitable and non-excitable tissues (Artigas & Gadsby, 2004; Kim et al., 1995; Muramatsu et al., 1988; Wang & Horisberger, 1997). The resultant 7.5 Å-wide pore can conduct more than a million cations per second as opposed to merely pumping a hundred cations per second (Artigas & Gadsby, 2004; Gadsby et al., 2009; Reyes & Gadsby, 2006). In this way, superfusion of 1 to 10 nanomolar PLTX elicits oxygen-glucose deprivation (OGD)-like SD in rodent brain slices (Kim et al., 2025), as well as initiating single or multiple SDs in the locust central nervous system (CNS) (Wang et al., 2024) as also recorded during locust anoxia. Several much earlier studies using brain slices have hinted that an SD activator might be released by energetically stressed tissue (see Discussion).

In the current study, we developed an experimental protocol to generate artificial cerebrospinal fluid (aCSF)-containing biomolecule(s) released from rat brain slices undergoing SD. We then used several independent techniques to partly characterize and isolate these biomolecules, which are hereafter referred to collectively as a putative spreading depolarization *activator*, SD*a* (Fig.1A,B).

**Figure 1.**
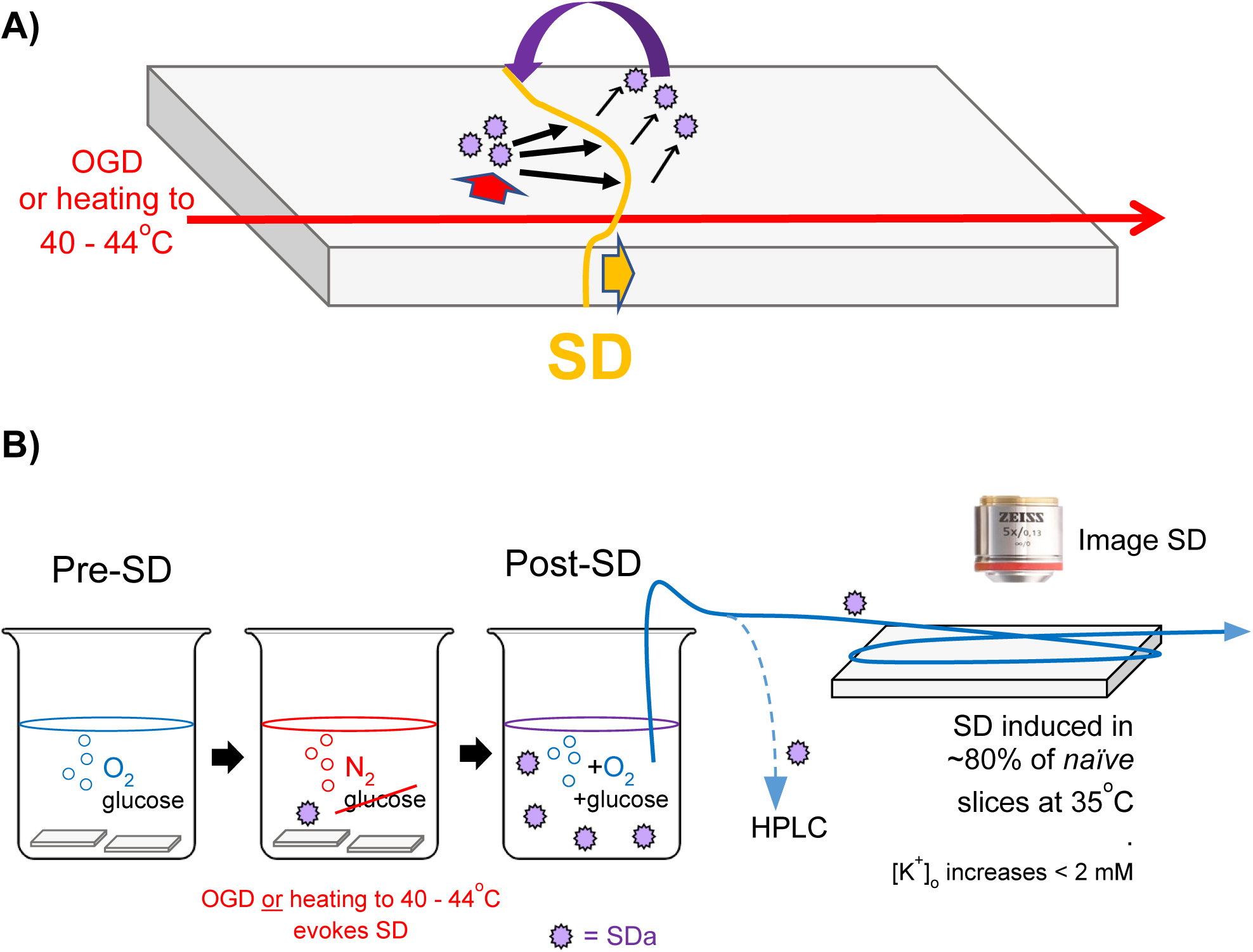
**A) Hypothesis:** An endogenous SD *activator* (SD*a, purple circles*), released during metabolic stress, is responsible for *initiating* (red arrowhead) and *promoting the propagation* of SD (yellow arrowhead). The self-regenerating feedback is represented by the curved arrow. **B) Testing the hypothesis**: SD evoked by O_2_/glucose derivation (OGD) or hyperthermia evokes SD, releasing an SD activator (circles) from slices which are then removed from the beaker. The post-SD aCSF is bioassayed for SD generation using a naive, healthy slice (right) or analyzed using high pressure liquid chromatography (HPLC).

Our results provide further support for our previous work implicating an SD*a* acting as a humoral factor released just prior to and during SD (Kim et al., 2025) (Fig. 1A,B). This then promotes regenerative SD, possibly by opening the NKA transporter in the well-documented PLTX-like manner.

## Materials and Methods

### Animals

All experiments were conducted under animal use protocols as approved by the Queen’s University Animal Care Committee based on standards set by the Canadian Council on Animal Care. Such protocols adhered to international standards of animal welfare to minimize the pain and suffering of animals. Male Sprague-Dawley rats (21 days) were obtained from Charles River Laboratories (Senneville, QC, Canada) for brain slice experiments and SD*a* sample preparation. Ages at experimentation were 3–6 weeks. For RBC experiments, male CD-1 mice (+22 days) from Charles River Laboratories were used. All animals were group-housed on a 12-hour light/dark cycle at 21°C (± 1°C) and were provided with rodent chow and reverse-osmosis water *ad libitum*.

### Experimental Solutions

Standard aCSF (pH ≈ 7.4) was prepared using double-distilled water (ddH_2_O) and the following concentrations of salts (in mM): 120 NaCl, 3.3 KCl, 26 NaHCO_3_, 1.3 MgSO_4_, 1.23 NaH_2_PO_4_, 11 D-glucose and 1.8 CaCl_2_. The aCSF was bubbled with 95% O_2_ / 5% CO_2_ throughout experiments. The exception was oxygen-glucose deprived aCSF (“OGD-aCSF”), prepared as described above but without glucose and instead bubbled with 95% N_2_ / 5% CO_2_ (Fig. 1B). Hyperthermic (Ht) aCSF was standard aCSF warmed from 35°C to 40-45°C and bubbled with 95% O_2_ / 5% CO_2_.

To prepare a PLTX stock solution, a single sample of 100 μg palytoxin (Wako Chemicals USA, Richmond, VA) was supplied on a clear film in a glass vial. To make a 10 μM stock solution, 3.73 mL of ddH2O was added to the vial which dissolved the film. This yielded about 3.7 mL of 10 μM palytoxin stock solution which was then serially diluted in aCSF for subthreshold PLTX experiments.

### Brain Slice Preparation

A rat was placed in a soft plastic DecapiCone^®^ (Braintree Scientific; Braintree, MA, USA) and decapitated using a guillotine. Brains were rapidly excised from the skull and flushed with ice-cold, oxygenated aCSF. The olfactory bulbs and cerebellum were removed. The brain was secured on a cutting platform against a small block of agar, bisected, submerged in ice-cold aCSF, and sectioned midline into 400 μm hemi-coronal slices using a Leica 1200-T vibratome (speed 0.09 mm/s; amplitude 0.70–0.85; temp 5-10° C; Leica Biosystems; Wetzlar, Germany). Each slice was gently transferred via wide-mouth pipette into an oxygenated recovery chamber containing aCSF at 35°C and incubated for at least one hour prior to use.

### Imaging Changes in Light Transmittance (ΔLT) Through Slices

Following one hour of recovery from slicing, a hemisected coronal rat brain slice was transferred to an imaging chamber on an inverted microscope (Fig. 1B) and superfused with gravity-fed solutions (2 mL/min) maintained at 35°C ± 0.5°C using a dual temperature controller (Warner Instruments; Hamden, CT, USA). The slice was transilluminated with a halogen light source using a 2.5X objective. Images were captured with a charge-coupled-device (Hamamatsu Photonics; Bridgewater, NJ, USA) and frame grabber (INDEC Biosystems; Santa Clara, CA, USA). Light transmittance (LT) through the tissue was digitized with frames acquired at 0.5 to 0.05 Hz using Imaging Workbench 6.0 software. Change in LT (ΔLT) was calculated according to the following equation:

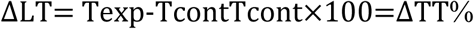

The first frame of the image series was the control (T_cont_) and was subtracted from each subsequent image (T_exp_). Because T_cont_ varies based on the zone sampled, the data were normalized by dividing by T_cont_. ΔLT could thus be expressed as a percentage of the control image intensity. Changes in LT were displayed using a pseudo-colored light intensity scale (low to high: blue-green-yellow-orange-red) where increased LT indicates tissue swelling. By contrast, reduced LT associated with dendritic beading was pseudo-colored purple (Fig. 2) (Andrew & MacVicar, 1994; Risher, W. C., Andrew, R. D., & Kirov, S. A., 2009).

**Figure 2.**
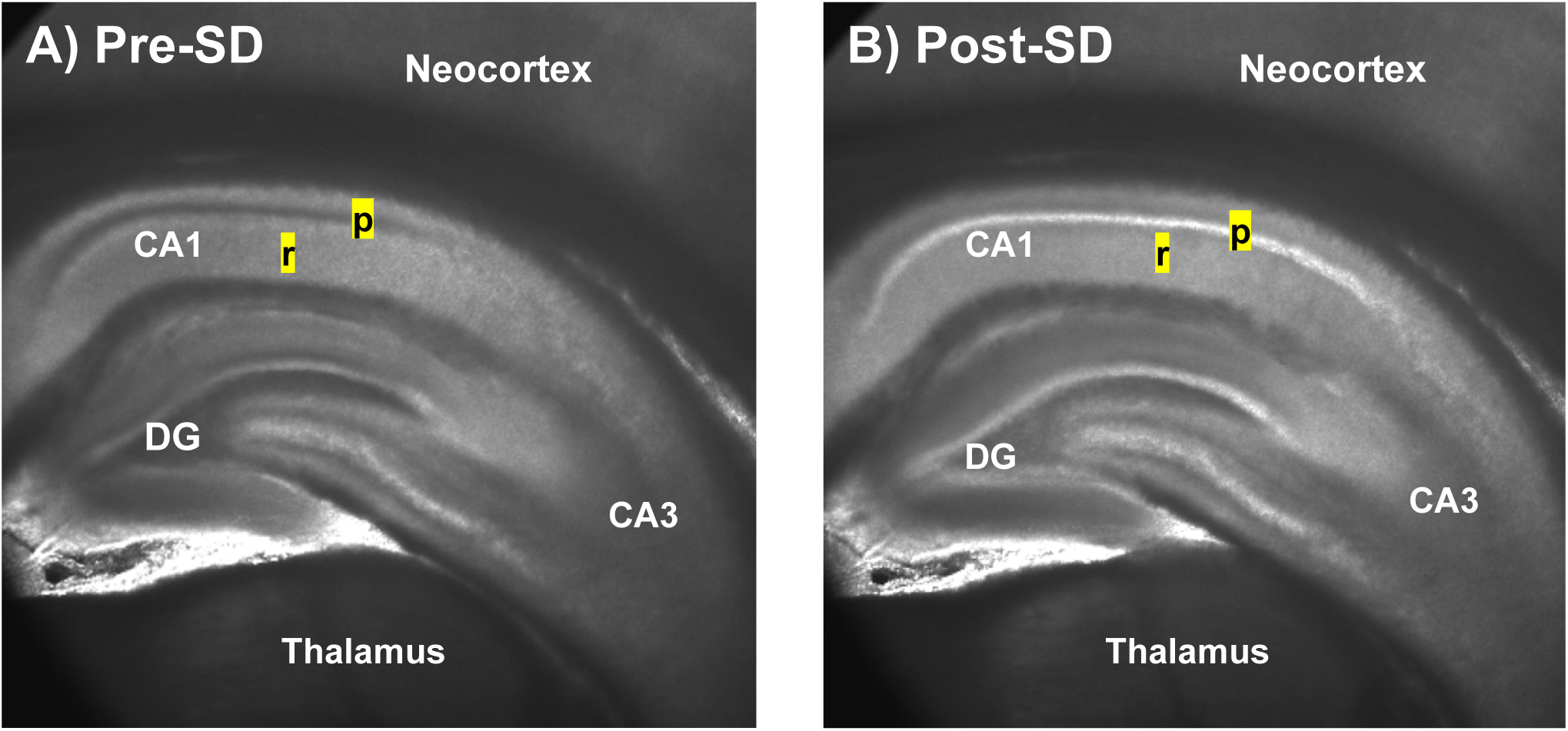
Simple light transmittance (LT) images reveal early SD injury. **A)** Before OGD-induced SD, the ‘dark’ CA1 cell body layer (stratum pyramidale, p) indicates a healthy hippocampus. **B)** Following SD, cell bodies swell, increasing the translucence of the stratum pyramidale in CA1 (p) and the granule cell body layer in the dentate gyrus (DG).

Specific LT imaging protocols are outlined in Table 1. **1)** Slices of neocortex with underlying hippocampus or striatum were imaged in flowing aCSF superfused for 2 minutes prior to application of test solutions, ensuring a stable baseline. **2)** To simulate ischemia, OGD-aCSF was superfused for 10 to 20 minutes following the baseline period. **3)** For PLTX threshold experiments, PLTX-aCSF was superfused at various concentrations for 10 minutes, followed by 10 minutes of OGD-aCSF superfusion to test if another SD could be evoked by OGD after PLTX application. This protocol was designed to identify the threshold [PLTX] that induced SD. **4)** For PLTX priming experiments, OGD-aCSF containing a subthreshold [PLTX] superfused slice. The control condition involved superfusion of aCSF alone for 20 minutes following baseline imaging. **5,6)**. For experiments testing for released SD*a,* 10 mL aliquots of Pre-SD or Post-SD_OGD_ (or Post-SD_Ht_) aCSF were thawed to room temperature. The freeze-thaw procedure did not alter the SD*a* activity, and occasionally fresh (non-frozen) aliquots were used. All 10 mL samples were then diluted with 35 mL aCSF. This increased the total volume of the superfusate, enabling longer image recordings lasting 10-20 minutes.

**Table 1.**
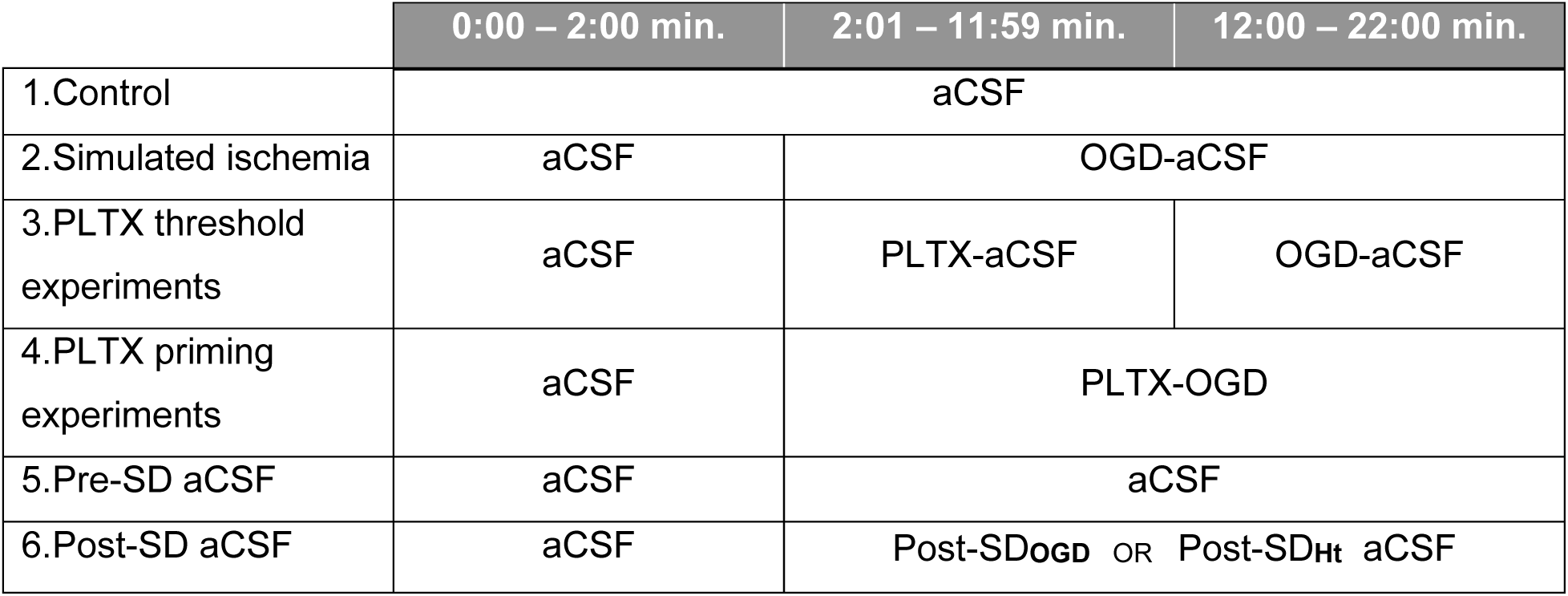
Superfusion protocols for brain slice experiments. After two minutes of baseline imaging in aCSF, brain slices were superfused with one of a variety of solutions while imaged for twenty minutes to monitor for SD onset. Beside each experimental condition (far left column) are the corresponding solutions used during each time period. For PLTX threshold and priming experiments, [PLTX] ranged from 0.01 to 1.0 nM.

|  | 0:00 – 2:00 min. | 2:01 – 11:59 min. | 12:00 – 22:00 min. |
| --- | --- | --- | --- |
| 1.Control | aCSF |  |  |
| 2.Simulated ischemia | aCSF | OGD-aCSF |  |
| 3.PLTX threshold experiments | aCSF | PLTX-aCSF | OGD-aCSF |
| 4.PLTX priming experiments | aCSF | PLTX-OGD |  |
| 5.Pre-SD aCSF | aCSF | aCSF |  |
| 6.Post-SD aCSF | aCSF | Post-SD<sub>OGD</sub> OR Post-SD<sub>Ht</sub> aCSF |  |

### Pre-SD and Post-SD_OGD_ aCSF Preparation

After recovery from dissection, 26–40 hemi-slices were transferred to a small mesh basket in the bottom of a 50 mL beaker containing standard aCSF bubbled with 95%O_2_ / 5% CO_2_ at 35°C (Fig. 1B). Next, the slice basket was transferred for 10 minutes to a new beaker containing 10 mL standard aCSF bubbled with 95%O_2_ / 5% CO_2_ at 35°C. This generated a “Pre-SD aCSF” as healthy slices do not undergo SD in standard aCSF at physiological temperatures. The basket was then transferred for 10 minutes to a beaker containing 10 mL of OGD-aCSF at 35°C <u>or</u> transferred to a beaker held at 40 to 44°C (hyperthermia, Ht). Exposure to either OGD or to Ht for this duration we found evokes SD in each brain slice. Either treatment generated “Post-SD aCSF”, which we further analyzed as described below.

Slice status after incubation in Pre-SD or Post-SD aCSF was occasionally confirmed using light microscopy. Specifically, a healthy hippocampus that has not undergone SD displays a lighter CA1 dendritic region (*stratum radiatum*, r) with a darker cell body layer (*stratum pyramidale*, p) (Fig. 3, left). In contrast, SD causes neuronal cell body swelling and dendritic beading, producing a more translucent cell body layer and darkened CA1 dendritic region (Fig. 3, right). Following a 10-minute incubation period, all Pre-SD and Post-SD solutions were filtered through a 40 μm cell strainer (Corning Life Sciences; Tewksbury, MA, USA) into 50 mL conical tubes. Solutions were aliquoted and then snap-frozen on dry ice, stored at -80°C prior to use, and used within 4 weeks of preparation.

**Figure 3.**
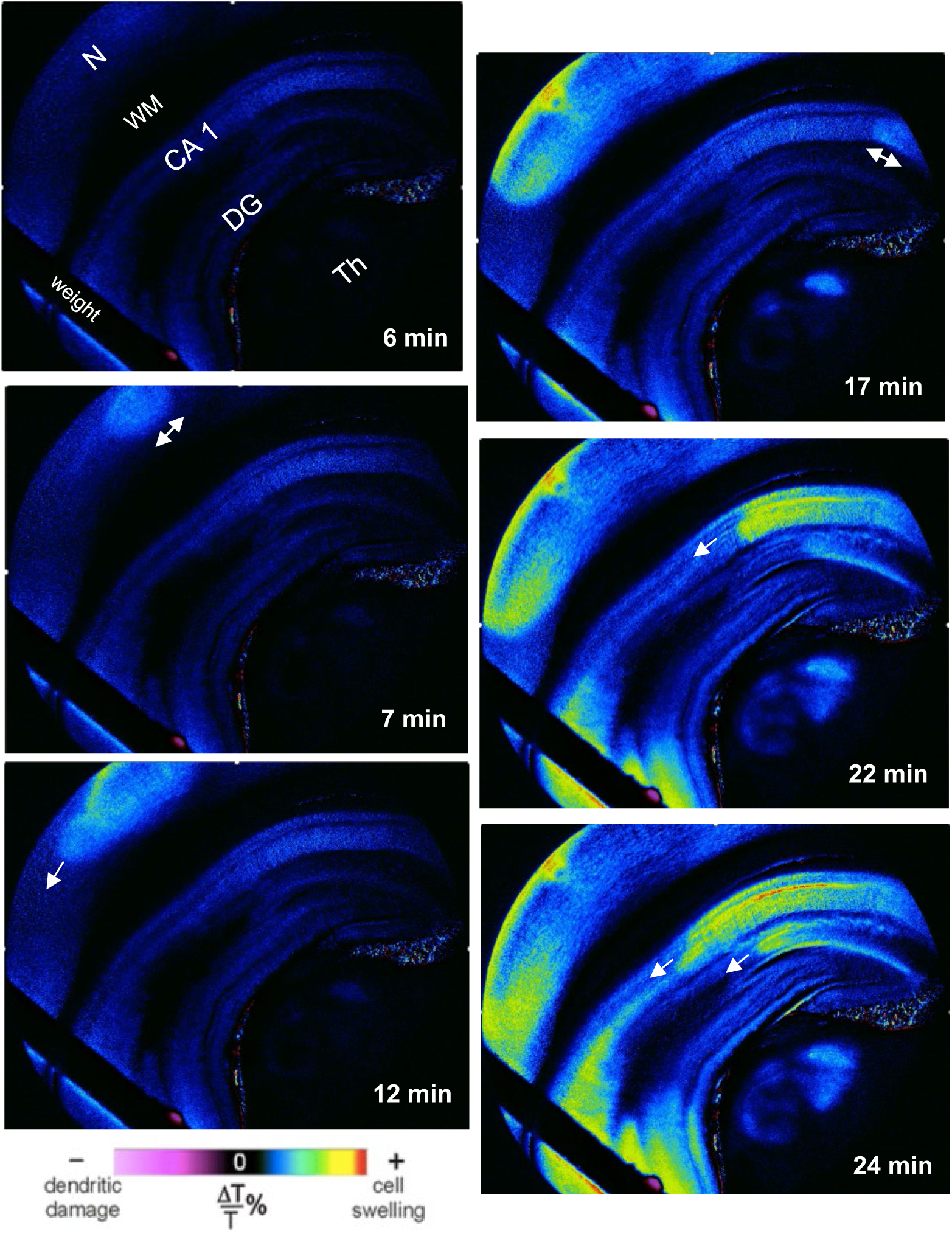
Coronal rat hemi-slice superfused with OGD aCSF at 35°C, evoking SD (arrows) in neocortex (N), CA1 pyramidal neurons (CA1), and dentate gyrus (DG). Thalamus (Th) showed some swelling but not white matter (WM). The imaging of this sequence did not extend long enough to detect dendritic damage.

### RBC Preparation

Whole blood was collected from CD-1 adult mice and immediately mixed with heparin (200 U/mL; 9:1 ratio of blood to anticoagulant). The blood was centrifuged at 300 x *g* for 10 minutes at 4°C. The supernatant (buffy coat and plasma) was removed, and RBCs were resuspended in 600 μL of 0.9% saline. The tube was gently inverted to mix, followed by centrifugation at 400 x *g* for 5 minutes. This wash step was repeated twice more, after which the supernatant was discarded. The packed RBCs were then resuspended in aCSF to create a 5% hematocrit stock for use in the hemolysis assay and the RBC swelling assay. This corresponded to a final reaction hematocrit of 0.25%, a concentration of cells at which the absorbance of the positive control (see below) fell within the range of linearity as per Beer’s Law (Mayerhöfer & Popp, 2019).

### Hemolysis Assay

The hemolysis assay was adapted from the protocol of Mazzarino et al. (2015). To generate a concentration-response curve for PLTX, experimental tubes contained varying volumes of vehicle (standard aCSF), 50 μL RBC stock, PLTX (0.01 to 0.5 nM) with or without ouabain (OUA, 5 to 500 μM) such that the final reaction volume was 1 mL. To assay SD samples, tubes contained 850 μL of either Pre-SD aCSF or Post-SD aCSF, 50 μL RBC stock, and in some cases, PLTX (0.005 to 0.05 nM). All tubes were brought up to a final volume of 1 mL using standard aCSF.

Positive controls (100% hemolysis) contained 50 μL RBC stock in 950 μL ddH_2_O; negative controls (0% hemolysis) contained 50 μL RBC stock in 950 μL standard aCSF. Tubes were incubated in a block heater at 37°C for 4 hours, with gentle inversion every 30 minutes to mix. All samples were assayed in duplicate. Following incubation, tubes were centrifuged at 500 x *g* for 10 minutes at room temperature to pellet intact RBCs. Finally, 200 μL of the supernatant was transferred to a clear, flat-bottomed 96-well microplate (Sarstedt; Nümbrecht, Germany) and the absorbance was measured at 540 nm using the SpectraMax iD3 Multi-Mode Microplate Reader (Molecular Devices; San Jose, CA, USA). Absorbance values for duplicates were averaged and the rate of hemolysis was calculated using the equation described by Mazzarino et al., (2015):

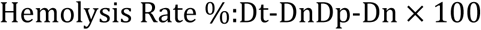

where *Dt*, *Dn*, and *Dp* are the average absorbances of the tested sample, negative control, and positive control, respectively.

To specifically monitor if elevated [K^+^]_o_ released by lysed RBCs could evoke hemolysis of RBCs, a separate hemolysis assay was performed in which RBCs were incubated in aCSF containing varying [K^+^]: 0, 1.1, 3.3 (standard aCSF), 5, 7, 15, and 26 mM. As in previous assays, 50 μL RBC stock was added to 950 μL aCSF.

### RBC Swelling Assay

We developed a more sensitive bioassay to detect early effects of Post-SD aCSF versus Pre-SD_OGD_ aCSF on RBCs. *Prior to* hemolysis, select hemolysis assays were repeated with corresponding light-microscopic visualization of RBC shape. At various timepoints (0, 15, and 30 minutes or 1, 1.5, 2, 3, and 4 hours) following the addition of the RBC stock to the assay tubes, 200 μL of reaction volume was removed from each assay tube and diluted 1:2 with standard aCSF. Next, 25 μL of the diluted sample was quickly pipetted onto a 12-well slide (MP Biomedicals; Santa Ana, CA, USA). RBCs were imaged with an inverted microscope, and a wide-field image of the cells was captured for later analysis. RBCs were counted and qualitatively classified as either biconcave (normal discoid shape, no swelling) or spheroid (swollen, loss of biconcave shape, smaller diameter) based on the parameters described in Carelli-Alinovi et al. (2014).

### [K^+^] Measurements

The [K^+^]_o_ in Pre-SD and Post-SD_OGD_ was measured using the ARCHITECT cSystem for clinical chemistry (Abbott Diagnostics; Chicago, IL, USA) by the core laboratory at Kingston Health Sciences Centre (Kingston, ON, Canada). Samples were kept on ice until measurement (∼ 2 hours).

### Statistical Analysis

Data were presented as mean (*M*) ± standard deviation (*StD*) and analyzed using a variety of statistical tests as appropriate, including unpaired Student’s *t*-tests and one-way ANOVAs with post-hoc Tukey tests. Statistical significance was set at *p* ≤ 0.05. Data were graphed and analyzed using GraphPad Prism^®^ 9 (GraphPad Software Inc.; La Jolla, CA, USA).

## Results

### Post-SD_OGD_ aCSF, but not Pre-SD aCSF, Evokes SD in Naïve Rat Brain Slices

To investigate whether a putative SD*a* solution is capable of evoking SD in a non-metabolically stressed rat brain slice, we superfused Pre-SD or Post-SD_OGD_ aCSF over a single healthy, naïve slice (Fig. 1B). Qualitatively, ΔLT imaging revealed no difference in slice health (i.e., no SD) after exposure to standard aCSF (data not shown) or to Pre-SD aCSF (Figs. 4A, 5A). Post-SD_OGD_ aCSF initiated *neocortical* SD in 82.35% of slices with an average propagation distance of 2.7 mm (Table 2; *n* = 17). This was true whether the Post-SD_OGD_ aCSF was used without freezing (Fig. 4B) or was frozen and thawed prior to use (Fig. 5B). The CA1 hippocampal region was comparatively resistant to SD evoked by Post-SD_OGD_ aCSF, with an SD frequency of only 5.9% (*n* = 17; data not shown). Thus, the CA1 region was not included in statistical analyses for SD onset time or for propagation distance. Overall, Post-SD_OGD_ aCSF reliably evoked SD in the neocortex of non-stressed slices, while Pre-SD aCSF did not.

**Figure 4.**
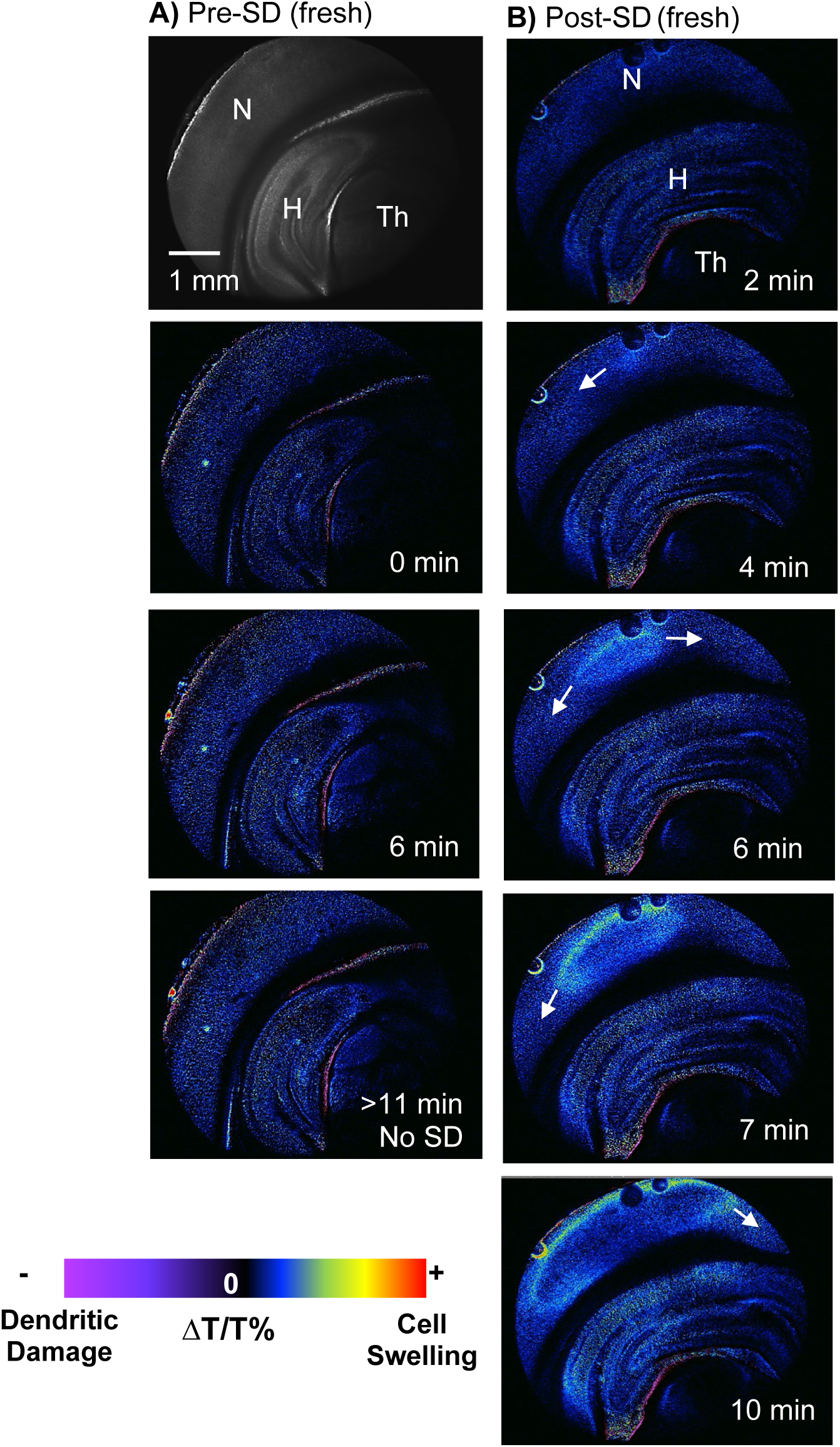
Naïve Rat Brain Slice Exposed to Freshly Prepared Pre- or Post-SD_OGD_ aCSF. The slices were superfused with either **A)** fresh Pre-SD aCSF (*n* = 8) or **B)** fresh Post-SD_OGD_ aCSF (*n* = 6). Only the latter caused SD (arrows). Time of image capture shown after the 2-minute baseline period. Neocortex (N), hippocampus (H), thalamus (Th), and direction of SD propagation (arrows) are indicated. The slices have not been stressed (e.g., by OGD or warming) in any other way.

**Figure 5.**
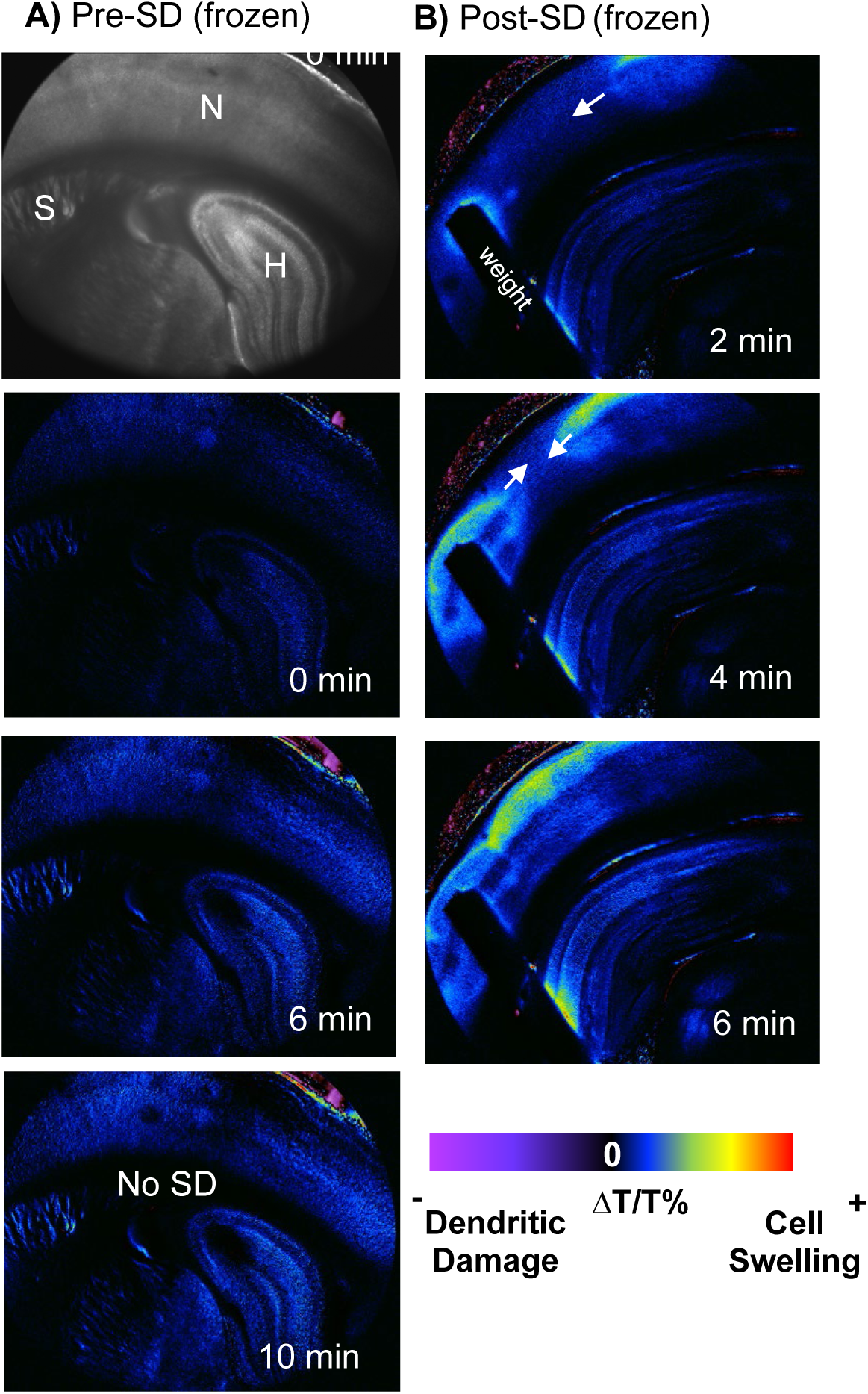
Rat Cerebral Slices Exposed to Frozen/Thawed Pre- or Post-SD_OGD_ aCSF. Rat cerebral slices superfused with either A) Pre-SD aCSF (*n* = 10) or B) Post-SD**_OGD_** aCSF (*n* =11), where both saline solutions were frozen to -80°C for at least 2 days. Time of image capture is after a 2-minute baseline period. Neocortical gray (N), hippocampus (H), striatum (S) and direction of SD propagation (arrows) are indicated. The slices were not stressed (by OGD or warming) in any way.

**Table 2.** Effects of Pre-SD aCSF versus Post-SD aCSF upon SD induction in the neocortex of naïve rat brain slices at 35°C. Pre-SD aCSF (*n* = 10) and then either Post-SD**_OGD_** aCSF (*n* = 17) or Post-SD**_Ht_** (n=23) was superfused over cerebral slices. Post-SD**_OGD_** aCSF or Post-SD**_Ht_** aCSF evoked SD in the neocortex while Pre-SD aCSF superfusion usually did not. * half of these slices had extracellular [K+] raised from 3.3 to 5.4 mM. a maneuver that did not increase SD frequency (not shown).

| Pre-SD aCSF |  |  |
| --- | --- | --- |
|  | No SD | SD |
| *n* (slices) | 10 | 0 |
| Average SD Distance (mm) | n/a | n/a |
| Post-SD aCSF |  |  |
|  | No SD | SD |
| Post-SD<sub>OGD</sub> : *n* (slices) | 3 | 14 |
| mean SD Distance (mm) | n/a | 2.7 |
| Post-SD<sub>Ht</sub> : *n* (slices) | 6/23 | 17/23 * |
| mean SD Distance (mm) | n/a | 2.5 |

To further quantify SD under the various experimental conditions, the following parameters were assessed: average SD onset time, average SD speed (mm/min), average ΔLT through the tissue, and maximum ΔLT through the tissue (Fig. 6). There was no significant difference in average onset time for neocortical SD evoked by Post-SD_OGD_ aCSF compared to OGD-aCSF alone (Fig. 6A). However, Post-SD_OGD_ aCSF-evoked SD propagating at a significantly slower speed as compared to OGD-evoked SD (Fig. 6B). Therefore, SD evoked by Post-SD_OGD_ aCSF in neocortex was similar to OGD-evoked SD in terms of onset time, but propagated more slowly.

**Figure 6.**
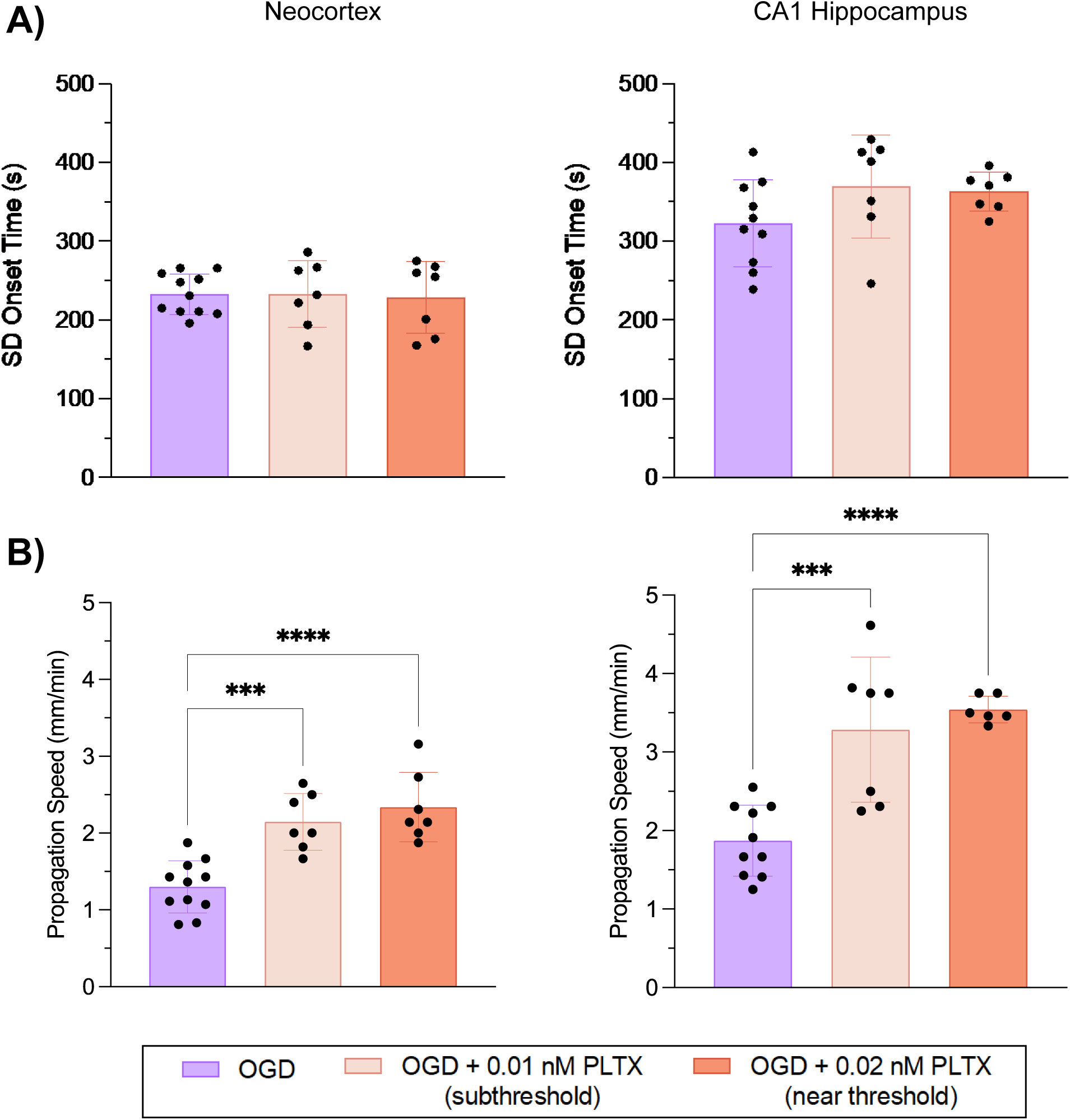
Subthreshold and near-threshold [PLTX] facilitates OGD-induced SD propagation speed, but not SD onset time. **A)** Average SD onset time (s) and **B)** SD propagation speed (mm/min) in the rat neocortex (left) and CA1 hippocampal region (right) following exposure to subthreshold PLTX-OGD (0.01 or 0.02 nM). For each experimental condition, *n* = 6 to 11. Each group was compared to OGD-aCSF (*n* = 10 to 11). Data are presented as mean ± *StD* and analyzed using a one-way ANOVA and a post hoc Tukey’s multiple comparisons test. (A, neocortex: *F*_2,22_ = 0.0294, *p* = 0.9711, *n* = 7–11; hippocampus: *F*_2,21_ = 2.107, *p* = 0.1466, *n* = 7–10; B, neocortex: *F*_2,22_ = 19.19, *p* < 0.0001, *n* = 7–11; hippocampus: *F*_2,20_ = 18.91, *p* < 0.0001, *n* = 6–10). *** *p* ≤ 0.001; **** *p* ≤ 0.0001.

Exposure to Post-SD_OGD_ aCSF significantly increased average neocortical LT change compared to standard aCSF and Pre-SD aCSF (Fig. 6C), but to a change similar to OGD-aCSF. Maximum ΔLT through the tissue is an indicator of the peak strength of SD at the wavefront. There was a significant effect of solution type on maximum ΔLT through the neocortex (Fig. 6D), such that Post-SD_OGD_ aCSF exposure significantly increased maximal neocortical ΔLT compared to standard aCSF and Pre-SD aCSF. Once again, application of Post-SD_OGD_ aCSF raised maximum ΔLT similar to OGD-aCSF. Taken together, these data show that Post-SD_OGD_ aCSF superfusion caused changes in LT consistent with SD generated by OGD. Importantly, Pre-SD aCSF did not elicit these changes because it could not evoke SD.

### PLTX Evokes Hemolysis of RBCs in a Concentration-Dependent Manner

To investigate the effect of Post-SD_OGD_ aCSF on the NKA more specifically, we used an RBC model, comparing hemolysis between Pre- and Post-SD_OGD_ aCSF, and a known NKA-specific inhibitor, PLTX. To start, an average concentration-response curve for the hemolytic effect of PLTX was generated (Fig. 7A; *n* = 3 assays per concentration). This confirmed that PLTX-evoked hemolysis of RBCs was concentration-dependent, with an EC_50_ of 0.0267 nM.

**Figure 7.**
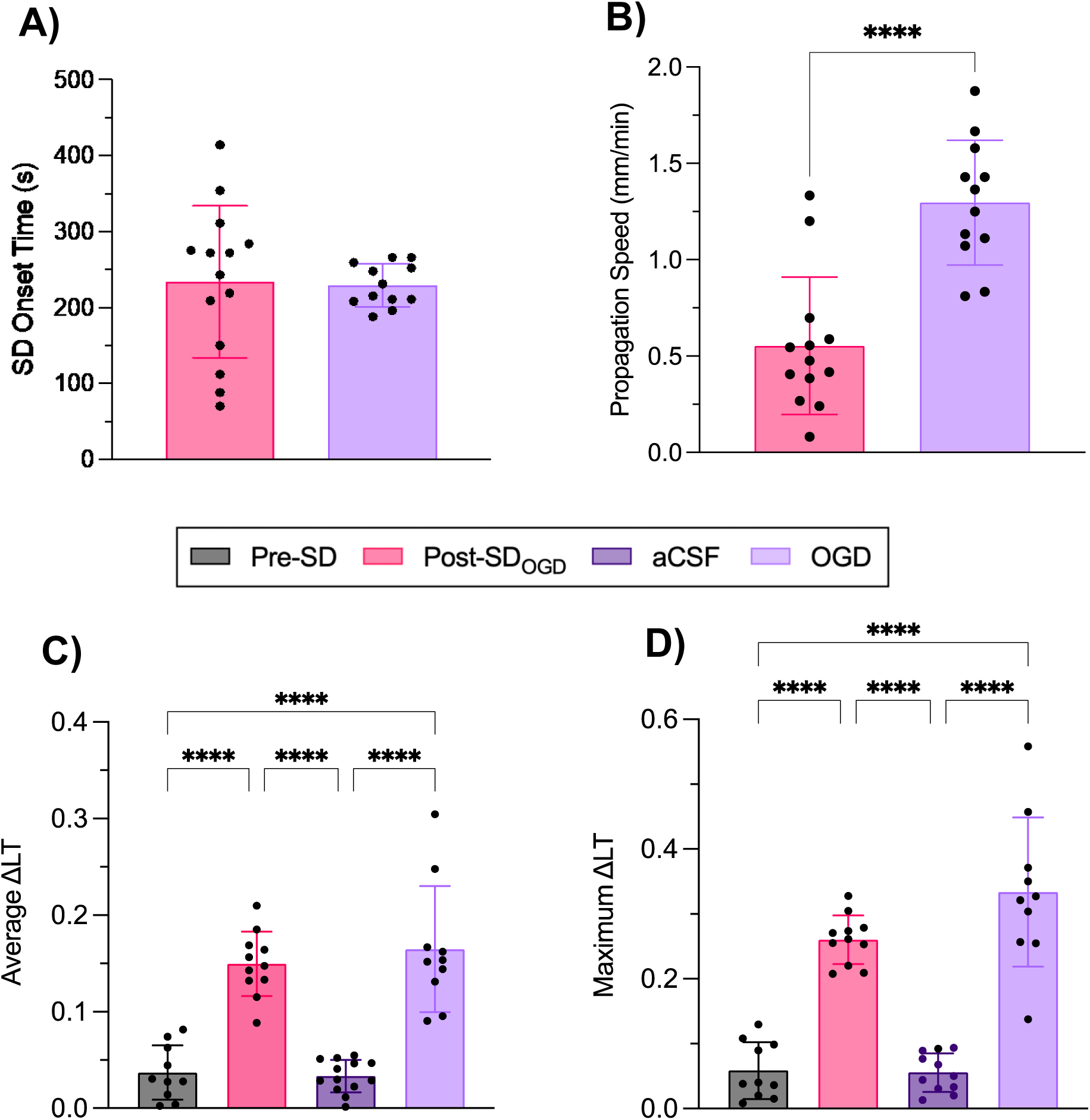
Effect of different salines on quantitative parameters of SD in the rat neocortex. **A)** Average SD onset time and **B)** average SD propagation speed from slices exposed to Post-SD**_OGD_** aCSF (*n* = 14) <u>or</u> OGD-aCSF (*n =* 12). Data are presented as mean ± *StD*. Statistical significance was determined using an unpaired Student’s *t*-test (A: *t*(24) = 0.1512, *p* = 0.8810; B: (*t*(23) = 5.432, *p* < 0.0001. Saline that bathed slices undergoing OGD displayed the same mean SD onset time as OGD itself but SD propagation speed was significantly slower. **C)** Average ΔLT and **D)** maximum ΔLT through the neocortex after superfusion of either Pre-SD aCSF (*n* = 10), Post-SD_OGD_ aCSF (*n* = 11), standard aCSF (*n* = 11–13), or OGD-aCSF (*n* = 10). ΔLT is measured as a change in LT from the baseline image in arbitrary units. Data are presented as mean ± *StD* and analyzed via one-way ANOVA and post hoc Tukey’s multiple comparisons test (C: *F_3,40_* = 36.38, *p* < 0.0001, *n* = 10–13; D: *F*_3,38_= 49.62, *p* < 0.0001, *n* = 10–11). **** *p* ≤ 0.0001. Both C and D show that LT increased significantly when SD was evoked.

Ouabain (OUA) is a known competitive inhibitor of PLTX as it binds to the NKA, so we investigated whether PLTX-induced lysis of RBCs could be attenuated by the addition of a fixed concentration of ouabain (250 μM; a concentration shown in a prior assay to reduce PLTX-induced hemolysis by >50% (Fig. 7C, blue). Consistent with previous assays by others, PLTX-evoked hemolysis was concentration-dependent (Fig. 7B; pink), with an EC_50_ of 0.0145 nM (95% *CI*: [0.0068, 0.1916]). The addition of a fixed concentration of OUA produced a rightward shift in the concentration-response curve (Fig. 7B; blue), increasing the EC_50_ of PLTX to 0.0428 nM. While this shows the expected attenuation of PLTX-induced hemolysis, OUA had no inhibitory effect at higher [PLTX] of 0.1 or 0.2 nM. This is in keeping with previous studies showing a higher affinity for NKA binding by PLTX vs OUA (see Discussion).

Ouabain itself had no hemolytic activity at increasing concentrations (Fig. 7C, green), but at 100 μM or more, OUA successfully inhibited the hemolytic effect of PLTX at 0.03 nM (Fig. 7C, blue). Further, when RBCs were pre-incubated with OUA for 30 minutes prior to PLTX exposure, or added to a solution of OUA and PLTX simultaneously, there was a similar inhibitory effect of PLTX-induced hemolysis (Fig. 7D).

### Post-SD_OGD_ aCSF Promotes PLTX-Induced Hemolysis of RBCs

Next, we tested whether Post-SD_OGD_ aCSF promoted hemolysis. Because Post-SD_OGD_ aCSF superfusion of a naïve brain slice induced SD while Pre-SD aCSF did not, we hypothesized that Post-SD_OGD_ aCSF contains the putative SD*a* and so would enhance hemolysis, if it were acting in a PLTX-like fashion. First, RBCs were incubated with Pre-SD or Post-SD_OGD_ aCSF. Surprisingly, there was no appreciable hemolysis in either group, suggesting that Post-SD_OGD_ aCSF alone does not induce RBC lysis. We then tested whether hemolysis could be provoked by Post-SD_OGD_ if the RBCs were *primed* by a very low concentration of PLTX which lacks hemolytic potency alone. To this end, we repeated the above hemolysis assays using Pre- and Post-SD_OGD_ aCSF with the addition of 0.005 to 0.05 nM PLTX (Fig. 8).

**Figure 8.**
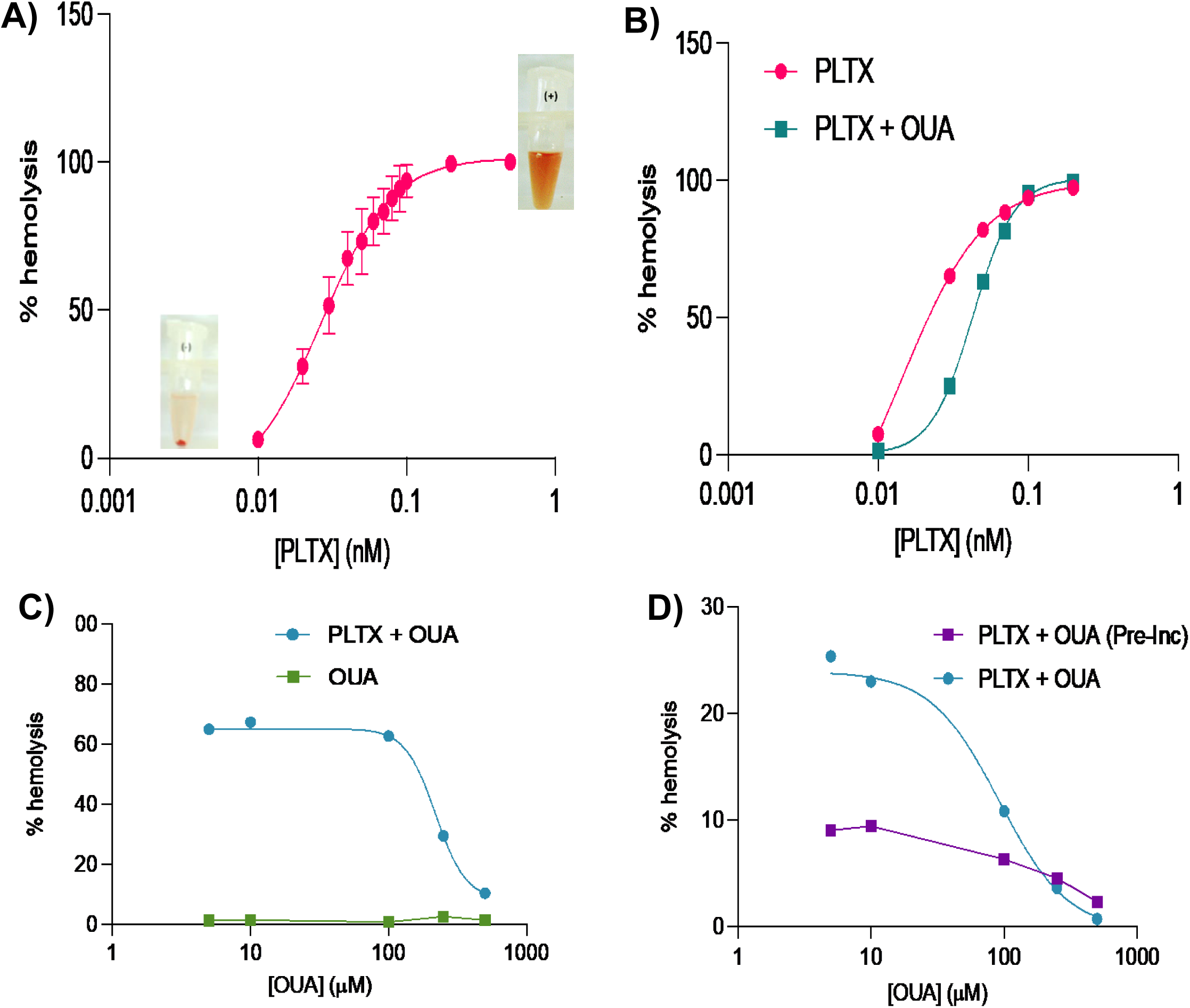
PLTX induces hemolysis of rat RBCs in a concentration-dependent manner which is inhibited by ouabain (OUA). **A)** RBCs incubated with [PLTX] = 0.01 to 0.5 nM. Data are mean ± *StD* (*n =* 3) and were fit to a non-linear regression model. **B)** Concentration-dependent hemolysis by PLTX is inhibited by 250 μM OUA. RBCs incubated with [PLTX] = 0.01–0.2 nM with or without 250 μM OUA. Data were fit to a non-linear regression model, revealing an EC_50_ for PLTX of 0.0145 nM (95% *CI*: [0.0068, 0.1916]) (red line). OUA induced a rightward shift in the concentration-response curve (green line), increasing the EC_50_ of PLTX to 0.0428 nM. **C)** Hemolysis of RBCs by PLTX is inhibited by OUA concentrations >100 μM. RBCs incubated with [OUA] = 5–500 μM with or without [PLTX] = 0.03 nM. **D)** Incubation with [PLTX] = 0.03 nM and [OUA] = 5–500 μM. RBCs were either pre-incubated with OUA for 30 minutes prior to the addition of PLTX (purple line) or added to OUA and PLTX simultaneously (blue line). OUA pre-incubation for 30 minutes potently inhibits PLTX-induced hemolysis. The blue line is fit to a non-linear regression model, revealing an IC_50_ for OUA of 92.35 μM. These data show that hemolysis of RBCs is mediated by the NKA.

The addition of 0.005 nM PLTX to either Pre-SD or Post-SD_OGD_ aCSF did not affect RBC hemolysis (data not shown). However, when 0.01 nM PLTX was added, a ‘priming’ effect was observed, such that significantly more hemolysis was induced by Post-SD_OGD_ aCSF + 0.01 nM PLTX versus Pre-SD aCSF + 0.01 nM PLTX (Fig. 8B; *n* = 4–8 per group).

A similar pattern was observed when the experiment was repeated using a PLTX concentration of 0.02 nM (Fig. 8C). Similarly, significantly more hemolysis resulted using RBCs exposed to Post-SD_OGD_ aCSF + 0.02 nM PLTX versus Pre-SD aCSF + 0.02 nM PLTX (*n* = 4–8 per group). When the PLTX concentration was increased to 0.05 nM, maximum hemolysis was observed in both conditions and there was no significant difference in hemolysis between groups (Fig. 8D, *n* = 4–8 per group).

Overall, the hemolysis assays were remarkably consistent, showing that Post-SD aCSF specifically displayed elevated SD*a* activity. At [PLTX] of 0.01 nM and 0.02 nM, Post-SD_OGD_ induced significantly more hemolysis compared to Pre-SD aCSF. This indicated a PLTX priming threshold between 0.005–0.01 nM with hemolytic saturation at less than 0.05 nM PLTX.

### Post-SD_OGD_ aCSF Promotes Early PLTX-Induced RBC Swelling

Given that the hemolysis assays suggested a PLTX priming threshold, we hypothesized that the putative SD*a* in the Post-SD_OGD_ aCSF may be acting on the RBC surface NKA in a PLTX-like manner to cause RBC swelling, but not lysis. To investigate the effects of Post-SD_OGD_ aCSF on RBCs *prior* to hemolysis, we characterized RBC morphology throughout the hemolysis assay. To do this, RBCs were incubated in the various aCSFs (either with or without PLTX), counted, and categorized as either spheroid (swollen) or biconcave (normal). Representative images at 2-hours and percent spheroid cells for each experimental condition are outlined in Figures 9 and 10, respectively.

**Figure 9.**
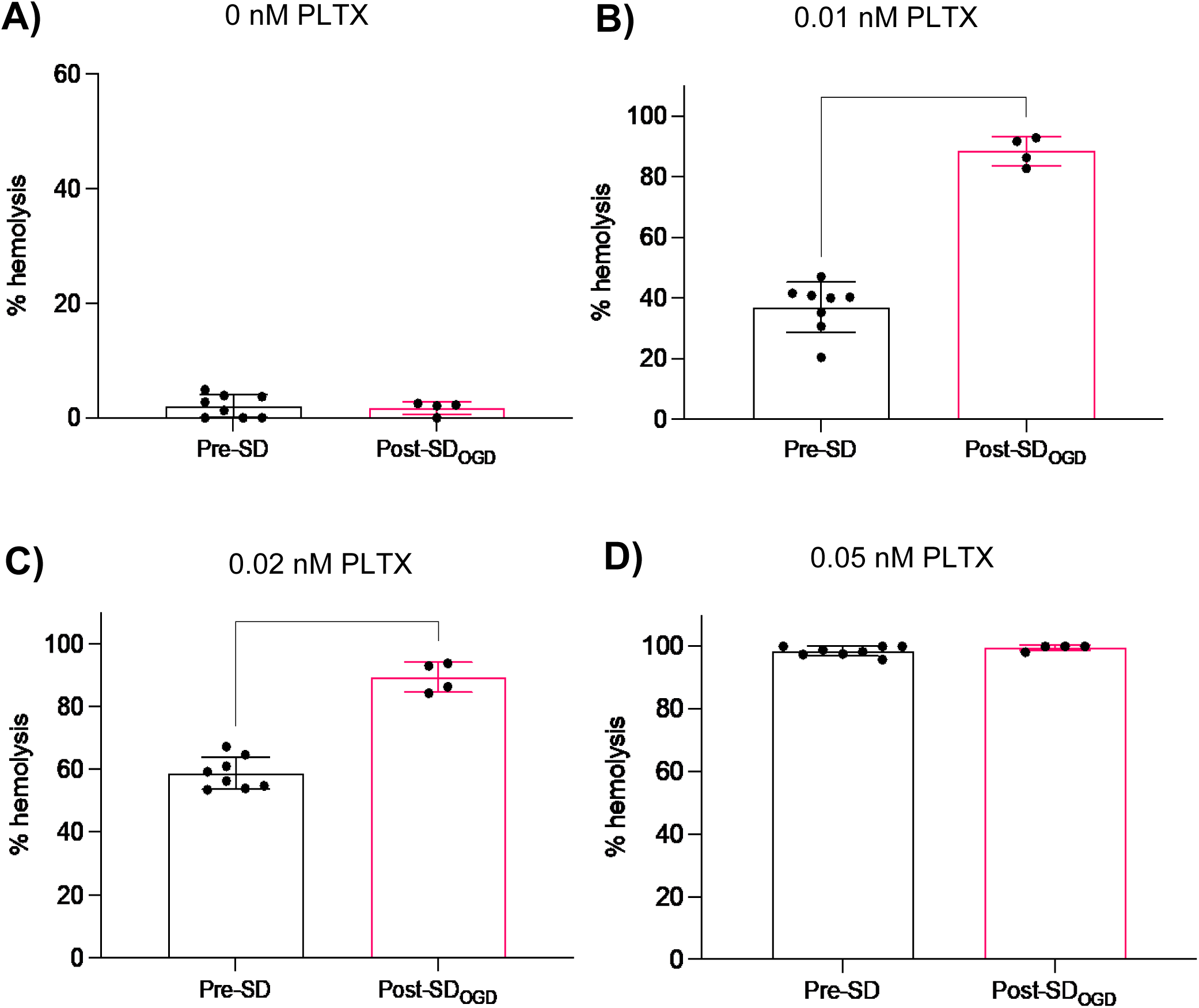
Post-SD_OGD_ aCSF potentiates PLTX-induced hemolysis of RBCs. The effect of Pre-SD aCSF versus Post-SD_OGD_ aCSF using different priming concentrations of PLTX on RBC hemolysis **A)** 0 nM PLTX shows no baseline hemolysis. **B,C)** 0.01 and.0.02 nM PLTX evokes baseline hemolysis (left, both histograms), and more hemolysis with the Post OGD saline right, both histograms. **D)** 0.05 nM PLTX saturates the responses. Data are presented as mean ± *StD* (*n =* 4–8) and analyzed by an unpaired Student’s *t*-test (A: *t*(10) = 0.3193, *p* = 0.7561; B: *t*(10) = 11.38, *p* < 0.0001; C: *t*(10) = 9.921, *p* < 0.0001; D: *t*(10) = 1.226, *p* = 0.2483). **** *p* ≤ 0.0001. Note that 0.01 nM PLTX <u>alone</u> in B has no hemolytic activity (see Figure 8A,B).

**Figure 10.**
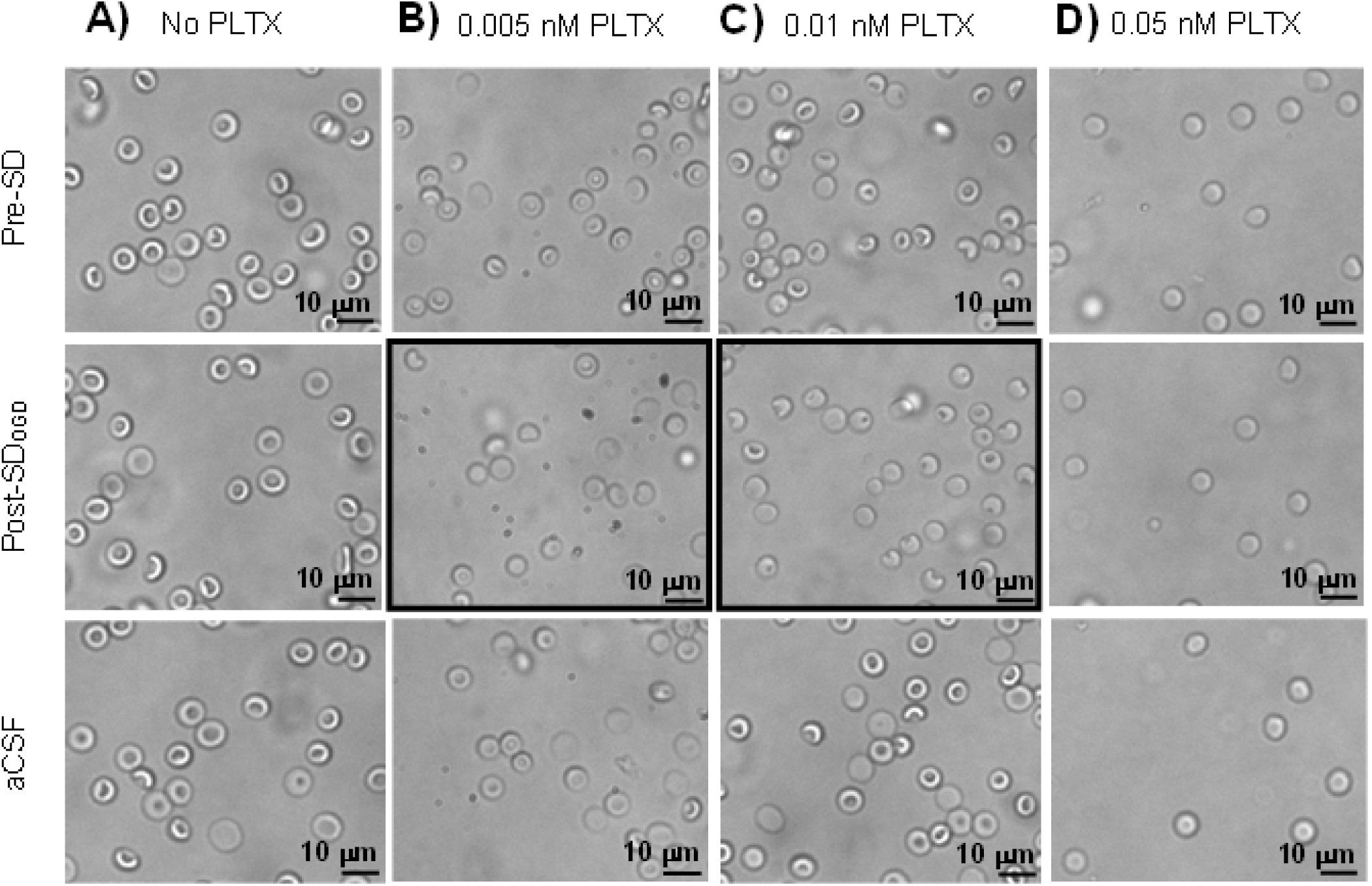
Representative images of RBCs under various incubation conditions mid-way through the swelling/hemolysis assay. Images show RBCs during a 2-hour incubation with Pre-SD aCSF, Post-SD_OGD_ aCSF, or standard aCSF at varying concentrations of PLTX. **A)** Zero PLTX shows no hemolysis, **B)** 0.005 nM PLTX and **C)** 0.01 nM PLTX show some return to biconcave shape after low [PLTX]. **D)** 0.05 nM PLTX evokes maximum hemolysis under each incubation. Note the greater swelling in Post-SD**_OGD_** aCSF (black boxes) compared to Pre-SD aCSF.

In the absence of PLTX, no RBC swelling was observed over 4 hours indicated by a consistent number of intact and biconcave RBCs in acquired images, regardless of incubation medium (Figs. 9A, 10A). This implies that Pre-SD aCSF, Post-SD_OGD_ aCSF, or standard aCSF did not induce RBC swelling alone.

When RBCs were incubated with 0.005 nM or 0.01 nM PLTX which were insufficient to induce hemolysis alone, swelling was observed between 1 and 1.5 hours in all conditions (Fig. 9B,C). The greatest swelling was observed in the Post-SD_OGD_ aCSF condition at every timepoint compared to both Pre-SD aCSF and standard aCSF (Fig. 10B,C). We interpreted this as elevated SD*a* activity in the Post-SD_OGD_ aSCF, where the NKA was ‘primed’ by the addition of a small amount of PLTX.

Finally, 0.05 nM PLTX rapidly swelled RBCs in all conditions, such that no differences between groups were evident (Figs. 9D, 10D). By 2 hours, all RBCs in all incubation conditions were swollen or lysed. As with the endpoint hemolysis assays, 0.05 nM PLTX-evoked swelling dominated the RBC response.

### [K^+^]_o_ is Only Slightly Elevated Post-SD

Release of extracellular K^+^ follows SD in brain slices. High enough [K^+^]_o_ (12 to 15 mM) can evoke SD. Therefore, to investigate if Post-SD_OGD_ aCSF contained enough [K^+^]_o_ to induce SD on its own, [K^+^]_o_ was measured. Post-SD_OGD_ aCSF had a significantly higher [K^+^]_o_ than Pre-SD aCSF (Fig. 11, *p* < 0.0001). Pre-SD aCSF had only slightly higher [K^+^]_o_ than control aCSF (no statistics performed due to *n* = 2 for aCSF). Thus, Post-SD_OGD_ aCSF which had superfused brain slices undergoing SD showed slightly elevated [K^+^]_o_ relative to Pre-SD aCSF, but not enough to induce SD alone.

**Figure 11.**
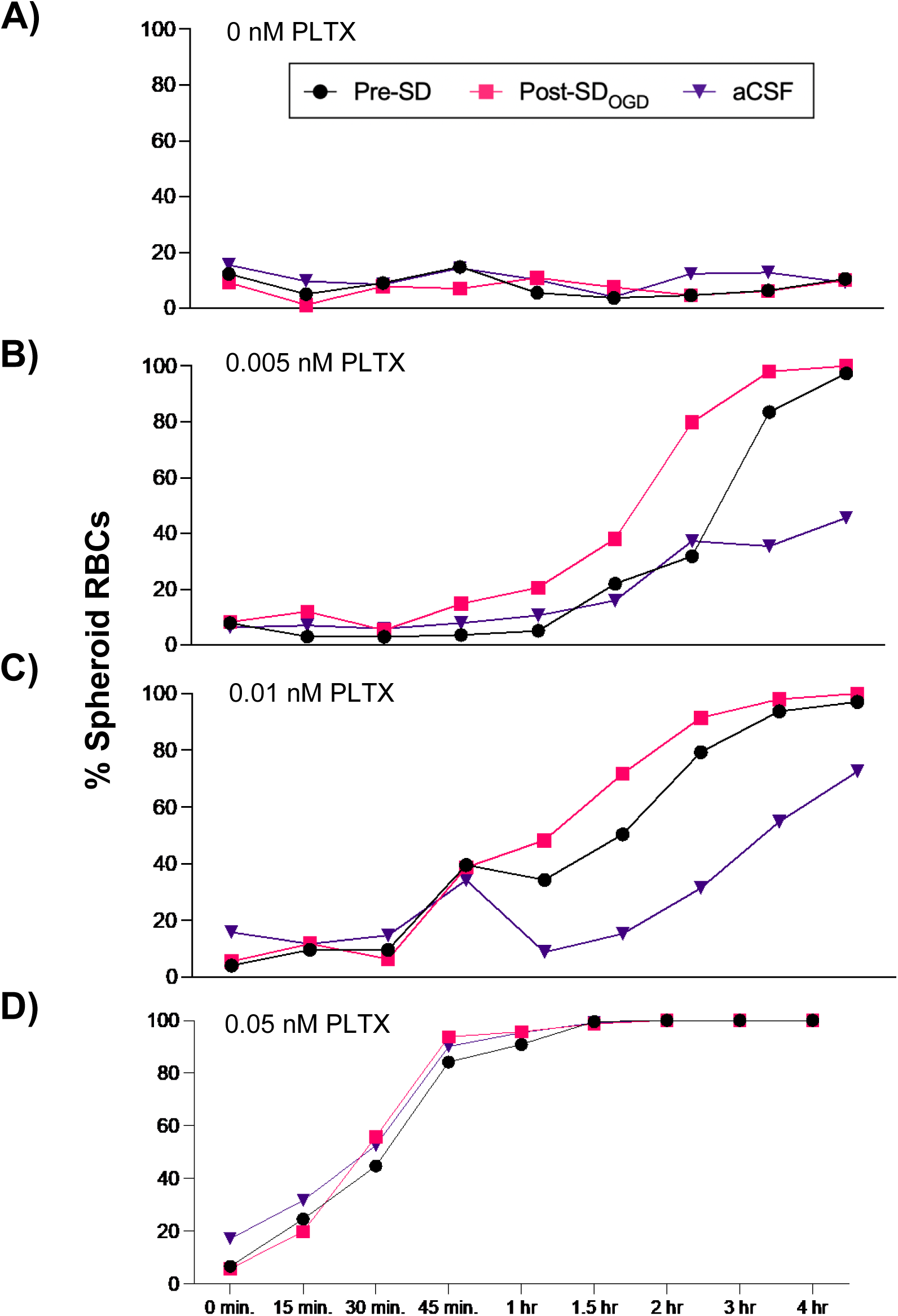
Time course of exposure to Pre-SD aCSF, Post-SD_OGD_ aCSF, or standard aCSF upon RBC swelling throughout the hemolysis assay. Y-axis shows the proportion of spheroid (swollen) RBCs under variable [PLTX] incubation conditions. **A)** Zero PLTX causes no swelling, **B)** 0.005 nM PLTX and **C)** 0.01 nM PLTX causes intermediate swelling, **D)** 0.05 nM PLTX evokes maximal swelling. Each data point represents a cell count from one wide-field image. Importantly, Post-SD**_OGD_** values are consistently greater than Pre-SD at 0.005 and 0.01 nM PLTX, both which are subthreshold concentrations for PLTX-evoked hemolysis alone (see Figure 8A,B).

### Elevated Extracellular [K^+^] Does Not Evoke Hemolysis of RBCs

To confirm the above finding, we investigated if the small increase in [K^+^]_o_ in Post-SD_OGD_ aCSF compared to Pre-SD aCSF could evoke hemolysis and thus contribute to the hemolytic assays. To this end, we incubated RBCs in standard aCSF solutions containing [K^+^]_o_ ranging from 0 to 26 mM. No hemolysis occurred at any [K^+^]_o_ tested (Table 3), indicating that slightly elevated [K^+^]_o_ was not a factor in the hemolytic effect.

**Table 3.** Changing extracellular [K^+^] has no effect on the hemolysis of RBCs. The extent of RBC hemolysis in response to increasing [K^+^] (0 to 26 mM) in aCSF. Hemolysis was calculated as per standard protocol and is relative to positive and negative controls (see Methods). * Physiological [K^+^].

| [K<sup>+</sup>] in aCSF (mM) | % Hemolysis |
| --- | --- |
| 26 | 0 |
| 15 | 0 |
| 7 | 0.1 |
| 5 | 0 |
| 3.3* | 0 |
| 1.1 | 0.1 |
| 0 | 0 |

### Do Subthreshold Concentrations of PLTX ‘Prime’ the NKA to Fail, Promoting SD_OGD_ ?

By converting the NKA into an open channel, PLTX evokes SD in brain slices. To determine if the PLTX ‘priming’ of RBC hemolysis could be replicated in live brain slices, PLTX-OGD aCSF was superfused onto adult rat brain slices and SD was monitored using LT imaging.

We determined the minimum [PLTX] required to initiate SD in adult rat brain slices to identify an appropriate ‘priming’ [PLTX] for SD in brain slices. PLTX-aCSF from 0.01 to 1 nM was superfused over healthy, naïve slices. Analysis of image recordings indicated that SD was generated at a threshold of 0.03 to 0.04 nM (Table 4). This compares to a threshold for evoking 100% RBC hemolysis (a much slower process) of ∼0.01 nM (Fig. 7B) and a threshold for eliciting RBC swelling of ∼0.005 nM (Fig. 9B).

**Table 4.**
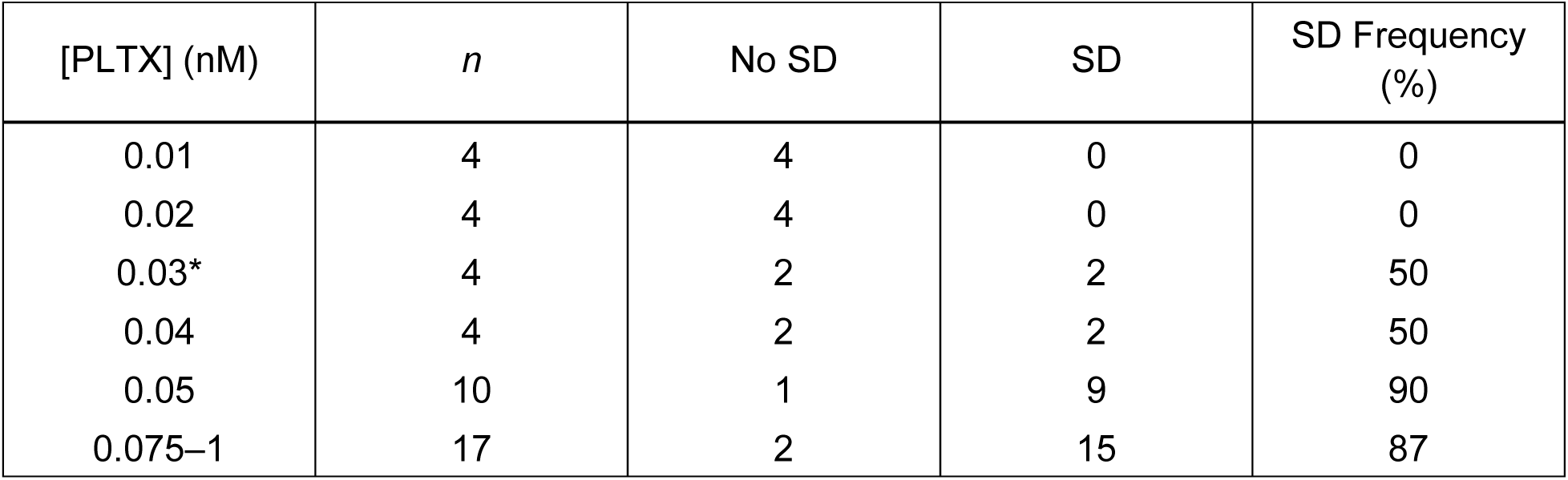
Determination of [PLTX] required for SD initiation in adult rat brain slices at 35°C. SD frequency was measured in response to superfusion of naïve rat brain slices with PLTX-aCSF (0.01–1 nM) (*n* = 4–10 per individual concentration). In general, SD frequency increased with increasing [PLTX]. *Approx. threshold of [PLTX] required to elicit SD in rat brain slices.

| [PLTX] (nM) | *n* | No SD | SD | SD Frequency (%) |
| --- | --- | --- | --- | --- |
| 0.01 | 4 | 4 | 0 | 0 |
| 0.02 | 4 | 4 | 0 | 0 |
| 0.03* | 4 | 2 | 2 | 50 |
| 0.04 | 4 | 2 | 2 | 50 |
| 0.05 | 10 | 1 | 9 | 90 |
| 0.075–1 | 17 | 2 | 15 | 87 |

Using 0.03-0.04 nM as the minimum [PLTX] required to provoke SD, we superfused OGD-aCSF containing subthreshold [PLTX] to assess for a ‘priming’ effect in brain slices. There was no priming effect for SD onset time in slices at low concentrations of PLTX (0.01 and 0.02 nM; Fig. 12A). However, with respect to SD propagation speed, bath application of 0.01 nM PLTX-OGD significantly increased speed in both neocortex (*p* = 0.0004) and hippocampus (*p =* 0.0003) relative to OGD-aCSF alone (Fig. 12B). Likewise, 0.02 nM PLTX-OGD increased propagation speed in neocortex (*p* < 0.0001) and hippocampus (*p* < 0.0001) compared to OGD-aCSF alone. Thus, subthreshold concentrations of PLTX that prime hemolysis of RBCs also speed up SD propagation in brain slices.

**Figure 12.**
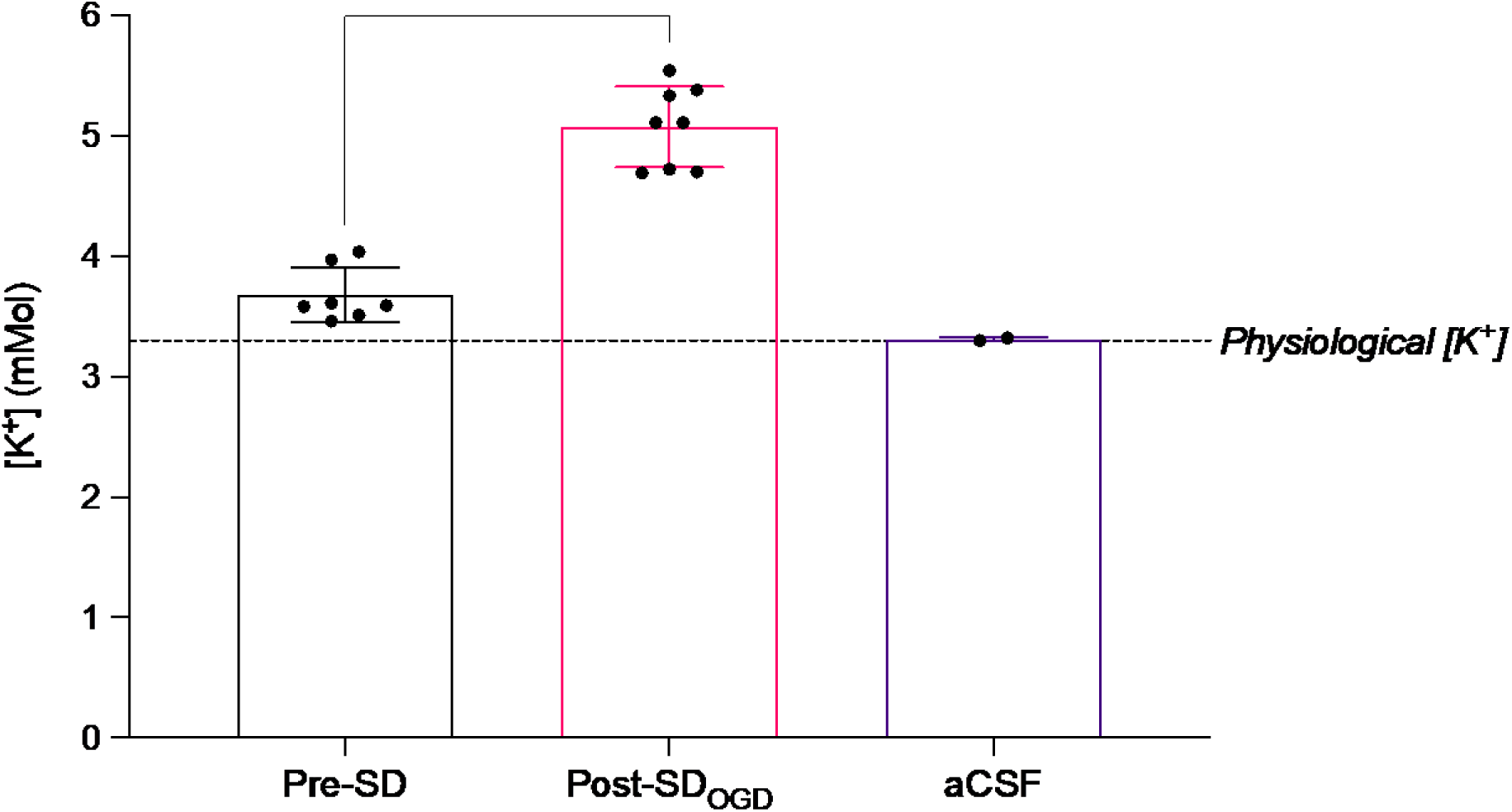
Release of extracellular [K^+^] increases by only 1.2 mM in Post-SD_OGD_ aCSF compared to Pre-SD aCSF. Thus increased [K^+^]_O_ is not the cause of SD in slices. Data are presented as mean ± *StD* (*n =* 7–8 per group) and analyzed by an unpaired Student’s *t*-test (*t*(13) = 9.246, *p* < 0.0001). **** *p* ≤ 0.0001. Slice-naïve standard aCSF (“aCSF”; *n* = 2) is shown for reference but was not included in statistical comparisons.

## Discussion

Recurring SDs arise in several neurological disorders and exacerbate the energy crisis in gray matter which is already metabolically compromised by ischemia or trauma. Each SD event expands the site of injury (Andrew, Farkas et al., 2022; Andrew, Hartings et al., 2022; Dreier, 2011; Pietrobon & Moskowitz, 2014) and worsens clinical outcomes (Chung et al., 2016; Lauritzen et al., 2011). However, little is known about the molecular mechanisms underlying SD initiation and propagation, particularly regarding which channel(s) open to drive the regenerative depolarization. Significantly, SD recurrence over the hours and days following the initial insult offers a therapeutic window during which SD could be inhibited to limit the overall extent of tissue damage (Andrew, Hartings et al., 2022; Hartings et al., 2003).

Both Leão (1944) and van Harreveld (1959) suggested that gray matter might release an endogenous compound that promotes or otherwise influences SD. Some scattered evidence over the past 50 years supports this idea. Martins-Ferreira et al. (1974) demonstrated that SD could be evoked in a naïve, untreated retina when the preparation was exposed to a medium that had been previously bathing a retinal preparation undergoing light-induced SD (Martins Ferreira et al., 1974). Kunkler and Kraig (1998) showed that hippocampal organ cultures, which do not typically undergo SD, become susceptible to SD by electrical stimulation. They hypothesized that soluble compounds released from the cells into the surrounding interstitium might be a requirement for SD. More recently, Kim et al. (2025) presented evidence that an endogenous molecule released during ischemia converts the NKA from a transporter to an open channel, thereby mimicking the well-documented action of PLTX.

The current study builds upon that work by investigating whether rodent gray matter stressed either by OGD or by warming acts to release an SD-promoting compound. We hypothesized that this putative ‘SD activator’ would both evoke SD in live brain slices as well as induce swelling/hemolysis of RBCs, both via opening of the NKA, as initially shown in RBCs by Habermann et al. (1981). Here we provide further evidence for a released endogenous SD*a* and propose that it has a PLTX-like action upon the NKA to generate and propagate SD.

### Post-SD_OGD_ aCSF, but not Pre-SD aCSF, induces OGD-like SD in Naïve Brain Slices

We first exposed naïve rodent brain slices containing neocortex and hippocampus to Pre-SD aCSF and then Post-SD_OGD_ aCSF. Superfusion with Post-SD_OGD_ aCSF induced neocortical SD in ∼83% of slices whereas Pre-SD superfusion did not evoke SD (Table 2). By contrast, the sliced hippocampal region was relatively resistant to this induced SD. The threshold for SD in the hippocampus in coronal slices is reported as higher than neocortex as seen with OGD-induced SD in coronal brain slices (Hellas & Andrew, 2021).

Onset times for this neocortical SD evoked by Post-SD_OGD_ aCSF versus standard OGD-aCSF were similar (Fig. 6A). The SD evoked by Post-SD_OGD_ aCSF qualitatively resembled the SD evoked by OGD-aCSF and demonstrated a similar LT increase at the SD wavefront (Fig. 6C). The SD induced by Post-SD_OGD_ aCSF propagated more slowly than OGD-induced SD (Fig. 6B). Importantly, Pre-SD aCSF was *incapable* of evoking SD, suggesting that Post-SD_OGD_ aCSF contains a substance that can produce OGD-like SD in non-metabolically stressed neocortical slices. This substance retains its activity when frozen and then thawed, if tested 1 to 2 days after preparation (Figs. 4, 5).

### PLTX Evokes RBC Hemolysis which is Attenuated by OUA

Once we confirmed that Post-SD_OGD_ aCSF, but not Pre-SD aCSF, could induce SD in otherwise healthy and naïve brain slices, we investigated the NKA as a possible target for the putative SD*a* by using an *in vitro* hemolysis assay. First, we characterized the concentration-dependent hemolytic effect of low concentrations of PLTX (0.01–0.5 nM) (Fig. 7A). The concentration-dependent hemolytic effect of PLTX was first described by Habermann et al. (1981) who showed that hemolysis occurred 1–2 hours after PLTX application and was preceded by an immediate K^+^ efflux from RBCs (Ahnert-Hilger et al., 1982; Habermann et al., 1981). They attributed the resultant decreased osmotic resistance of RBCs to PLTX’s disruption of cell membrane permeability through an unknown mechanism (Ahnert-Hilger et al., 1982). Today, it is understood that PLTX acts by binding extracellularly to the NKA and converting this transporter into a non-specific cation channel (Hirsh & Wu, 1997; Scheiner-Bobis et al., 1994; Kim et al., 2025). In support, we confirmed that OUA which is a competitive inhibitor of PLTX can attenuate PLTX-evoked hemolysis up to a point. Inhibition with 250 μM OUA only occurred at PLTX concentrations less than 0.1 nM. Higher [PLTX] out-competes OUA for NKA binding (Wu, 2009).

### Very low [PLTX] ‘Primes’ the NKA of RBCs for SDa-induced RBC Swelling and Hemolysis

We next investigated whether exposure of RBCs in Post-SD aCSF evoked swelling, similar to how superfusion of Post-SD_OGD_ aCSF induced SD in live rat brain slices. Exposure to Post-SD_OGD_ aCSF alone did not induce hemolysis, but there was a striking priming effect when RBCs were incubated in Post-SD_OGD_ aCSF with very low concentrations of PLTX (Fig. 8). Significantly, more hemolysis was evoked in tubes containing Post-SD_OGD_ compared to Pre-SD aCSF, suggesting that there is a substance within the Post-SD_OGD_ aCSF that acts on the RBCs in a similar way to PLTX. The NKA is ‘primed’ for opening by a small concentration of PLTX. This effect cannot be explained by exposing the solution to brain slices, because Pre-SD aCSF had significantly less hemolytic potency when combined with the same concentration of PLTX.

However, because Post-SD_OGD_ aCSF did not induce hemolysis on its own, we investigated pre-lytic swelling of RBCs with the addition of a small concentration of PLTX, to detect any ‘priming’ effect (Figs. 9, 10). Again, there was a ‘priming’ effect of pre-lytic RBC swelling of the Post-SD_OGD_ aCSF with the addition of a small amount of PLTX insufficient to induce hemolysis on its own. The proportion of spheroid RBCs were higher in the Post-SD_OGD_ aCSF compared to Pre-SD aCSF, or to aCSF alone, suggesting a molecule within the Post-SD_OGD_ aCSF that is capable of causing RBC swelling in a PLTX-like manner.

### Facilitation of SD in Brain Slices and Potentiation of PLTX-induced Hemolysis by Post-SD aCSF is not caused by increased [K^+^]_o_

We next measured [K^+^]_o_ in Pre- and Post-SD_OGD_ aCSF. As expected, we found that the [K^+^]_o_ was slightly elevated in Post-SD_OGD_ aCSF relative to Pre-SD aCSF. Extracellular K^+^ release is a known consequence of SD (Vyskočil et al., 1972), so these findings support that the slices did in fact undergo SD, while slices incubated in Pre-SD aCSF did not. Notably, the average [K^+^]_o_ in Post-SD_OGD_ aCSF was far below the 12 mM threshold required to induce SD in brain slices (Ayata & Lauritzen, 2015).

### Subthreshold [PLTX] ‘Primes’ the NKA for OGD-induced SD in Naïve Slices

To investigate whether PLTX has a ‘priming’ effect on induction of SD in brain slices, we first determined the minimum [PLTX] required to elicit SD in live brain slices to be 0.03-0.04 nM (Table 4). Then, we exposed slices to OGD-aCSF containing subthreshold concentrations of PLTX (Fig. 12). SD propagation speed was increased with the addition of subthreshold PLTX (0.01–0.02 nM), in a concentration-dependent manner in both neocortex and CA1 hippocampus. This suggests a similar ‘priming’ effect with small concentrations of PLTX and the putative SD*a* released from the metabolically stressed live brain tissue undergoing OGD.

### Conclusions and Future Directions

Failure of the NKA with consequent disruption of Na^+^ and K^+^ gradients are the critical events initiating SD. An intriguing question is why Post-SD_OGD_ aCSF could elicit SD in brain slices, but did not evoke hemolysis of RBCs on its own. In RBCs, small “priming” amounts of the NKA inhibitor PLTX were required to initiate swelling and eventual hemolysis evoked by Post-SD_OGD_ aCSF containing the putative SD*a*. Undoubtedly, there is a reduced metabolic demand of an RBC compared to a neuron, the result of very low NKA density within the RBC plasma membrane (Gatto & Milanick, 2009). The high metabolic demand by neurons because of their high NKA density (Pivovarov et al., 2019) likely render neurons much more susceptible to NKA inhibition.

There were no differences using either OGD or hyperthermia (Ht) as the trigger of SD generation. Both initiated SD and showed identical HPLC traces of released molecules (see Appendix). Taken together, these findings provide support for the existence of an SD*a* released from metabolically-stressed gray matter which is then capable of inducing SD in unstressed brain slices. Also, the SD*a* elicits ouabain-sensitive RBC swelling and eventual hemolysis with the addition of small concentrations of PLTX which ‘prime’ its activity. It is possible that the very low density of RBC surface NKAs compared to neurons (Gatto & Milanick, 2009) means that RBCs require a minor ‘boost’ by trace amounts of PLTX to begin offsetting the transmembrane Na^+^ and K^+^ gradients. We also demonstrate that in live brain slice models, solutions containing the SD*a* are capable of provoking SD in naïve brain slice tissue, and a similar ‘priming’ phenomenon is seen with the addition of subthreshold amounts of PLTX, suggesting a small synergistic effect between the SD*a* and PLTX.

Future studies using HPLC and mass spectroscopy should attempt to purify solutions containing the SD*a* to enable a more robust characterization of this molecule’s structure. Ultimately, an improved understanding of the molecular events underlying SD initiation and propagation will offer novel therapeutic targets for reducing recurrent SD events to improve clinical outcomes of stroke, traumatic brain injury, and global brain ischemia.

## Abbreviations

aCSF: Artificial cerebrospinal fluid
AMPA: α-3-hydroxy-5-methyl-4-isoxazolepropionic acid
ATP: Adenosine triphosphate
AU: Arbitrary units
CNS: Central nervous system
ddH_2_O: Double-distilled water
HPLC: High-performance liquid chromatography
LT: Light transmittance
*Δ*LT: Change in light transmittance
M: Mean
NMDA: *N*-methyl-D-aspartate
OGD: Oxygen-glucose deprivation
OUA: Ouabain
PLTX: Palytoxin
RBCs: Red blood cells
SD: Spreading depolarization
SD*a*: Spreading depolarization activator
*StD*: Standard deviation

## Appendix

### High-Performance Liquid Chromatography (HPLC)

We used HPLC to analyze Pre-SD and Post-SD aCSF to detect and semi-quantitate released biomolecules from incubated brain slices. The primary goal was to demonstrate an increased release of biomolecules post-SD rather than characterizing the molecules themselves.

Filtered superfusate was injected into the HPLC column. To approximate their molecular weight range, GPC-100 columns (Eprogen/Promigen Life Sciences; Downers Grove, IL, USA) were roughly calibrated with known globular protein standards (not shown). GPC-PEP columns, which are ideal for small peptides (0.8–30 kDa), were calibrated with known linear peptide markers (not shown), all dissolved in ddH_2_O. The GPC-100 column was specific for both linear (0.5–25 kDa) and globular (5–160 kDa) molecules and can separate out water-soluble polymers such as proteins, nucleic acids, and carbohydrates. Thus 100 μL of either Pre-SD or Post-SD aCSF (100 μL samples) were analyzed using a Shimadzu LC-10AD HPLC apparatus (Shimadzu; Kyoto, Japan), equipped with either GPC-100 or GPC-PEP size-exclusion columns (Eprogen/Promigen Life Sciences; Downers Grove, IL, USA) with a mobile phase of 1X phosphate-buffered saline. Columns were calibrated with known linear peptide markers and proteins/peptides detected by fluorescence (excitation wavelength of 275–280 nm, emission wavelength 305 nm). Fluorescence (arbitrary units, AU) was normalized against the mobile phase for each run and subsequently plotted against elution volume using the Class-VP 7.2.1 SP1 software (Shimadzu; Kyoto, Japan). The molecular weight range of unknown biomolecules were then estimated by comparing their elution volumes with those of known protein/peptide standards (not shown).

### Size-Exclusion HPLC Analysis of Pre-SD and Post-SD aCSF

To search for further evidence of biomolecular release during SD and to approximate their size ranges, we used size-exclusion HPLC to analyze Pre-SD aCSF and Post-SD aCSF. Resultant peak patterns were characterized based on the linear peptide standards run on the GPC-PEP column (not shown). Peak amplitudes of the Pre-SD HPLC trace were assigned as 100%. Peaks from the Post-SD_OGD_ sample were compared as a percentage relative to the Pre-SD peak using an unpaired Student’s *t*-test.

In both Pre-SD and Post-SD_OGD_ aCSF, a predominant peak cluster was noted between elution volumes of 1.0 to 2.7 mL, roughly corresponding to a molecular weight range of 60 kDa (Fig. 14A1, ‘peak 1 cluster’; *n* = 8). Single peaks at elution volumes of 3.3 and 4.1 mL, roughly correspond to molecular weights between 2 kDa and 0.6 kDa respectively (Fig. 14A1, ‘peak 2’ and ‘peak 3’, respectively). Importantly all peaks, particularly peak 2, around 2 kD were significantly greater in Post-SD_OGD_ aCSF (n=8) compared to Pre-SD aCSF (*n* = 8).

**Figure 13.**
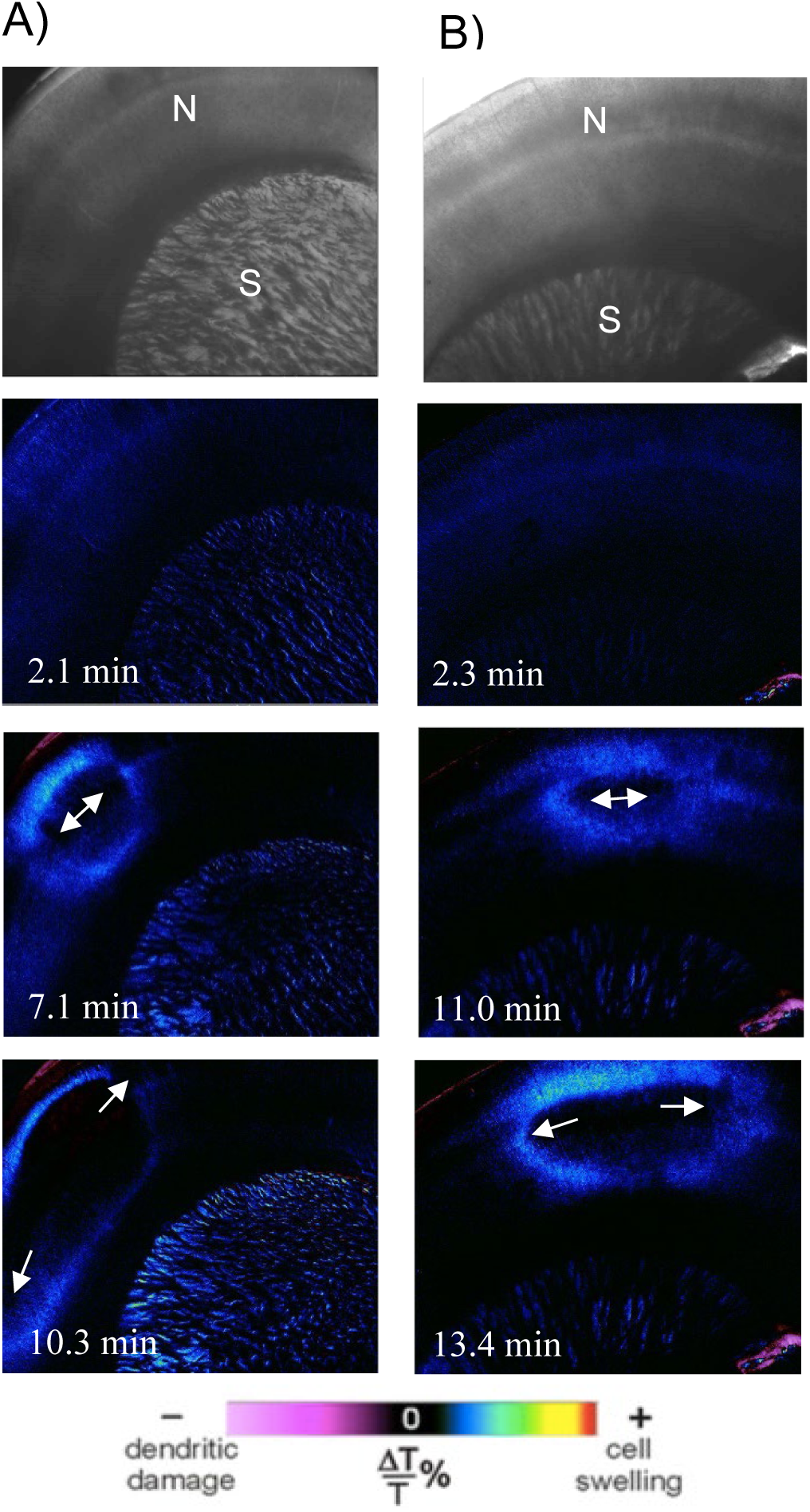
**A)** Abrupt hyperthermia (Ht), where slice temperature is raised from 35°C to 42°C over 5 minutes, initiates SD_Ht_ propagation (arrows) in rat neocortex. **B)** When the aCSF *bathing* such slices post-SD (as in A) is then superfused at 35°C over a *naïve* healthy slice, SD is evoked (as in B). The slice in B has not been warmed nor stressed in any other way. Neocortex (N); striatum (S).

**Figure 14.**
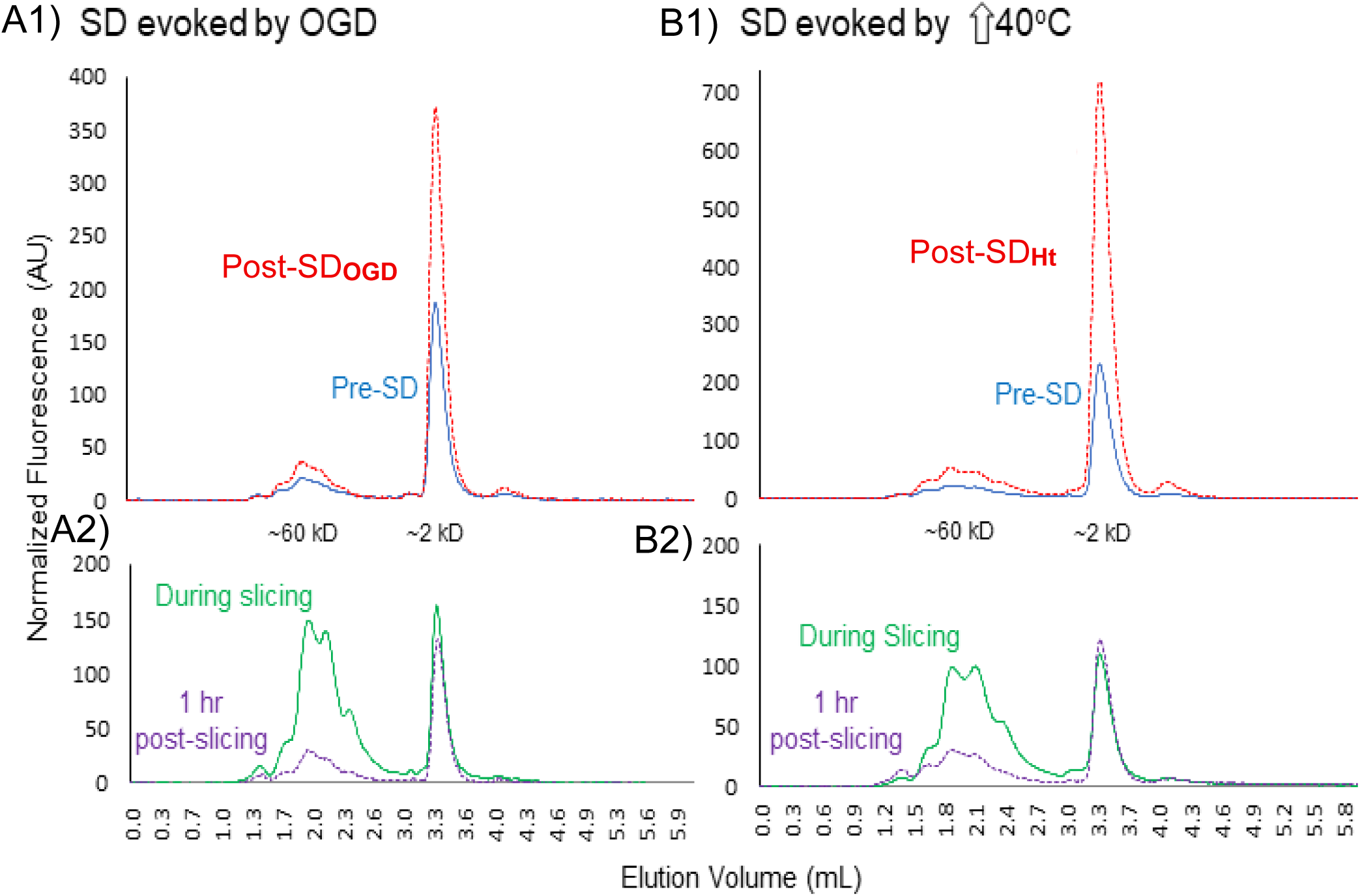
Post-SD aCSF contains more fluorescent molecules than Pre-SD aCSF (using a GPC-100 HPLC column) following either OGD-induced (left) or heat-induced (right) SD. Averaged traces of **A1)** Pre-SD aCSF versus Post-SD_OGD_ aCSF, and **B1)** Pre-SD aCSF versus Post-SD_Ht_ aCSF. **A2 and B2**) Standard aCSF bathing brain slices during and immediately following slicing. ACSF collected during actual slicing is associated with high levels of the fastest eluting biomolecules (green traces) likely caused by cell damage. Much lower levels are seen 1 hr later (purple traces) after slice recovery. Data analyzed by an unpaired Student’s *t*-test. * *p* ≤ 0.05, **** *p* ≤ 0.0001.

There is the potential confound that tissue slicing could cause nonspecific compound release into the aCSF because of cell damage/rupture at the slice’s cut surfaces. To address this, the aCSF “During Slicing” over 10 minutes was compared to aCSF bathing the slices for 10 minutes *after* slicing (at the end of the standard 1-hour incubation) which we termed “1-hr Post-Slicing” (Fig. 14A2, B2; *n* = 5 animals each). For analysis, peaks on the “1-hr Post-Slicing” HPLC trace were assigned as 100%, and peaks from the “During Slicing” trace were compared as a percentage using an unpaired Student’s *t*-test. The same peak 1 cluster as well as peak 2 were observed in both samples. The peak 1 cluster was significantly greater in the “During Slicing” samples relative to “1 hr Post Slicing” *n* = 3). There was no significant difference in peak 2 during vs post-slicing (Fig. 14A2) (*n* = 3). Thus., it appears that compounds in only the peak 1 cluster are associated with general tissue injury resulting from the slicing process. HPLC analysis of standard aCSF not exposed to slices yielded no peaks (data not shown).

Notably, all the HPLC peaks released from slices exposed to OGD for 10 minutes were remarkably similar to those elicited when SD was evoked by hyperthermia (Ht) specifically by raising slice temperature from 35°C to 40-44°C for 10 minutes (compare Figs.14B1 and B2). This shows that the molecules outside the peak 1 cluster were released by the process of SD and not tissue damage from slicing.

## Acknowledgements

The authors thank Dr. Susan Boehnke for her assistance with the HPLC and related data. CAL developed the hemolysis assay and performed and analyzed all experiments involving RBCs. JAH performed the *Δ*LT imaging experiments, analyzed the associated data, and prepared corresponding figures. RDA and NOB developed the initial methodology for generating Pre-SD and Post-SD aCSF, which CAL refined. NOB performed HPLC experiments and analyzed the associated data. CAL researched and wrote the manuscript. JAH and RDA assisted with writing and editing the manuscript. All authors assisted in revising the manuscript.

